# Hierarchical replay of multi-regional sequential spiking associated with working memory

**DOI:** 10.1101/2023.10.08.561458

**Authors:** Ermeng Huang, Da Xu, Huangao Zhu, Zhaoqin Chen, Yulei Chen, Xiaoxing Zhang, Chengyu T. Li

**Author notes:** These authors contributed equally (Ermeng Huang, Da Xu, Huangao Zhu).

## Abstract

How does millisecond-scale neural activity mediate behaviors over seconds? We recorded brain-wide activity in mice performing an olfactory working-memory task to decipher cross-region organization of activity. Spike-correlograms revealed millisecond within- and cross-region spike couplings, more prominent among neurons encoding similar memories. Spike coupling linked neurons into motifs of chains, single loops, and nested loops, especially among hippocampal and prefrontal-cortex neurons. Direction of spike coupling and activity chains was in line with that of memory-associated activity waves. Intriguingly, activity motifs were replayed before and after task performance, and during inter-trial intervals. Motifs were hierarchically organized, with progressively increasing time constants and the number of participating neurons. Thus, hierarchically organized and replayed cross-region spiking motifs are modulated on demand during delay period to mediate perceptual working memory.

**One Sentence Summary:** Nested activity motifs of chains, single loops, and nested loops, with progressively increasing time constants and number of participating neurons, are hierarchically organized and replayed to mediate perceptual working-memory maintenance.

## Main Text

The brain operates across a spectrum of temporal scales (*1*), spanning from the rapid millisecond-scale spiking activity to more prolonged second-scale behaviors like working memory (WM) (*2–4*). Working memory involves the active retention of both external and internal information, mediated by delay-period activity seen in various regions (*5–20*). A central question of systems neuroscience is how neuronal spiking is organized across temporal scales to mediate such behaviors.

Many theories have been proposed to solve the problem. The cell-assembly hypothesis (*21–23*) posits that sequential spike activity within a network characterized by recurrent connections could outlast short-lived external inputs (depicted in Fig. 1A). Building on the importance of recurrent connectivity, the attractor network theory (*18, 24–26*) underscores the sustained neuronal firing (as illustrated in Fig. 1B). Also depends on arrangement of connectivity, the synfire chain model (*27–29*) suggests the importance of the sequential propagation of spikes through pools of concurrently active neurons in a feed-forward manner (as shown in Fig. 1C). Additionally, the concept of a locally unstable yet globally stable trajectory of population dynamics (*9, 15, 30*) could potentially underlie the cross-temporal coordination of neuronal activity (depicted in Fig. 1D).

**Fig. 1.**
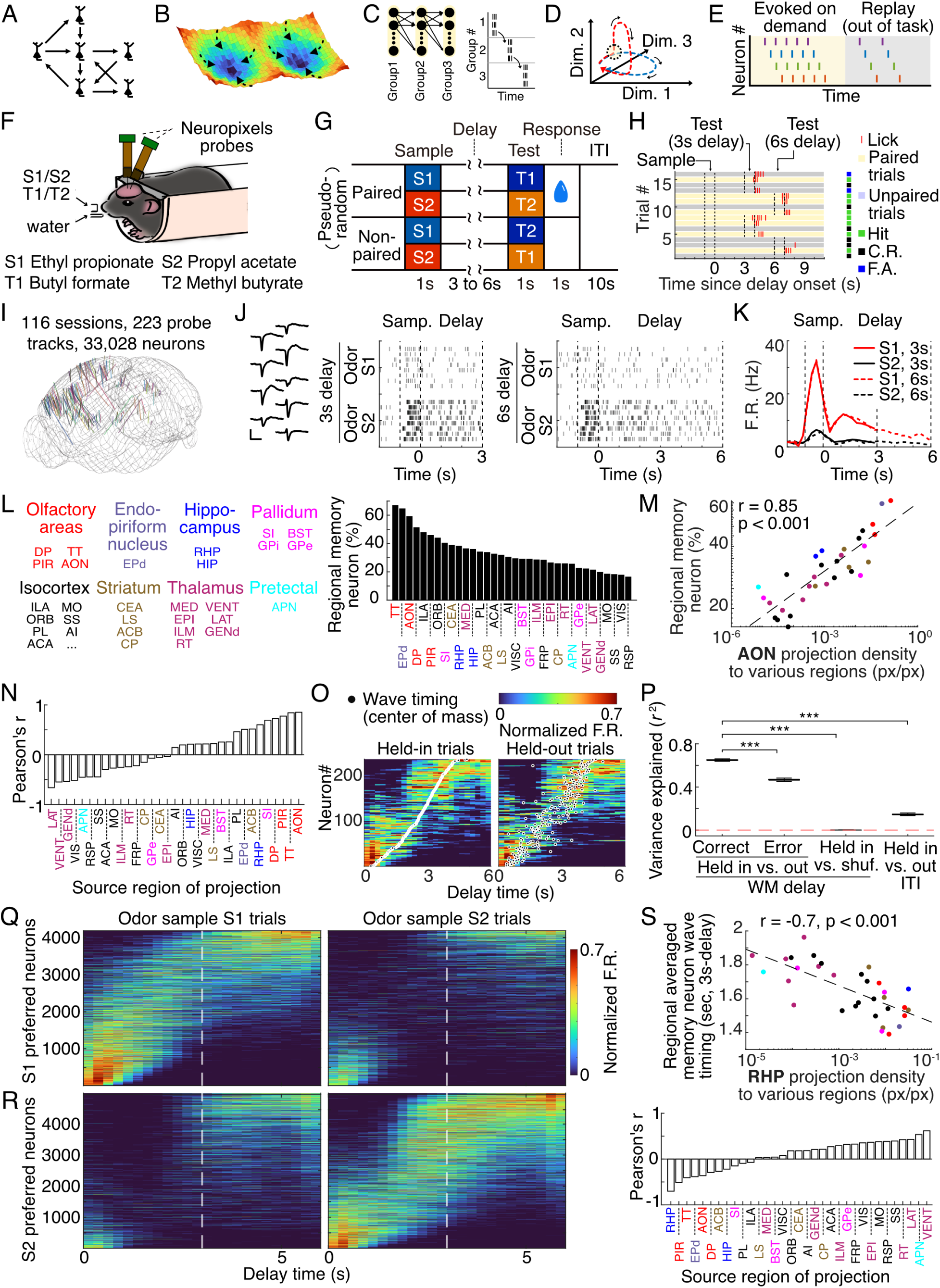
Brain-wide patterns of neuronal activity associated with WM-information maintenance. (**A-D**) Illustration of different theories of neuronal-spiking organization: cell assembly (A), point attractor dynamics (B), synfire chain (C), and transient population trajectory (D). (**E**) Illustration of neuronal spikes on demand and in replay. (**F**) Schematics of the behavioral training and recording setup. Below: odors used in the task. (**G**) Design of the olfactory delayed pair-association (ODPA) task. Sample, sample-odor delivery. Delay, delay period. Test, test-odor delivery. Response, response window. ITI, inter-trial interval. (**H**) Performance for exemplary consecutive trials in the task. (**I**) Co-registration of 223 probe tracks from 113 recording sessions on 40 mice into the Allen common reference space. (**J**) Activity of an example memory neuron (defined by the neurons with statistically significant different firing between two sample odors during the delay period). Left: Spike waveform on 9 consecutive recording sites. Scale bars represent 1 msec and 20 mV. Middle: Spike-raster plots of 8 example trials for two sample odors under 3-sec delay duration. Right: Spike raster for 6-sec delay duration. (**K**) Peristimulus time histogram (PSTH) of the example neuron in (J). (**L**) Distribution of the proportion of memory neurons (total number of neurons: 24,667) across 34 brain regions (only regions with more than 100 neurons registered are shown). Text color indicates high-level groupings of brain regions throughout Figure 1. Refer to table S1 for full name and the recorded neuron count of each region. (**M**) Correlation between regional proportions of memory neurons and projection density from AON to each of these regions, using data from the Allen Brain Atlas, r and p values represent Pearson’s correlation coefficient. (**N**) Distribution of correlation between the regional proportions of memory neurons and projection density from recorded regions throughout the brain. (**O**) Activity wave of an example recording session. Left: Odor-specific activity wave revealed by sorting the memory neurons by temporal center-of-mass (TCOM, red circles) from half of the correct trials (held-in). Right: Validation of activity wave for the other held-out trials, neuron ID as in the held-in trials. (**P**) For WM delay: Variance explained (r-squared) between TCOM of held-in trials and that of held-out trials, error trials, and shuffled held-out trials during the delay period for all recorded sessions (n = 116). For ITI: Variance explained (r-squared) between TCOM of held-in and that of held-out trials during the ITI. ***, p < 0.001, Wilcoxon rank-sum test. Whiskers, full range. (**Q**) Activity waves for the memory neurons that prefer sample S1 (n = 4,186). The order of the neurons was determined by TCOM. (**R**) As in (Q), for the memory neurons that prefer sample S2 (n = 4,996). (**S**) Relationship between regional wave timing and anatomy. Top: Correlation between regional averaged wave timing of memory neurons and projection density from RHP to each of these regions, using data from the Allen Brain Atlas. Below: Distribution of correlation coefficients between regional averaged wave-timing of memory neurons and output-projection density for the recorded regions.

In experimental observations, delay-period activity within different regions displays a spectrum of dynamic patterns. These patterns span from sustained firing resembling a point-attractor state (*5, 10, 18, 19, 25, 26*) to transient trajectories characterized by population waves and sporadic individual firing events (*6, 7, 9, 15, 16, 30*). Because that neural circuits extending across multiple brain regions collectively contribute to behavior (*12, 20, 31*), brain-wide mechanisms become necessary for understanding the organizing principles of neuronal firing underlying behavior.

What is the nature of temporal specificity for the global neuronal dynamics that underlie WM? Such activities could be exclusively triggered by the demand for WM maintenance (as depicted in Fig. 1E, left), or they might be spontaneously replayed beyond the delay period, for instance, during the inter-trial intervals or before and after behavioral sessions (as shown in Fig. 1E, right). Activity replay involves the reactivation of neural firing patterns that originally occurred during specific experiences or learning instances, largely accompanied with slow-wave ripple and with time compression of past experience (*32–34*). This intriguing neural phenomenon was observed during both sleep (*35, 36*) and wakefulness (*37–39*), prominently in the hippocampus and neocortex (*32–41*).

Replay could enable the offline reactivation and consolidation of memories (*32–34*), facilitate working memory of sequentially presented items (*42*), prevent catastrophic forgetting in continuous learning (*43*), and provide a mechanism to build internal representations that improve learning and decision-making (*44, 45*). Whether and how working memory-related spiking activity could be replayed outside of the delay period remains unclear.

Recent advances in Neuropixels-probe recordings (*46*) allow simultaneous recording of spikes from hundreds of neurons in many regions of behaving animals, uncovering brain-wide neural correlates of behavior at millisecond resolution (*47–52*). In the current study, we used dual Neuropixels probes to record spiking activity of more than 33,000 neurons in 62 brain regions in mice performing an olfactory WM task. By analyzing the dynamic structures linked by correlated spiking, we found that nested activity motifs of loops and chains are hierarchically organized during the delay period to support WM-related activity waves. Such activity motifs were replayed outside of the delay period and predominantly involved the hippocampal and prefrontal cortical neurons.

## Results

### Olfactory WM task and Neuropixels recordings

We trained mice to perform an olfactory paired-association (ODPA) task. Head-fixed mice (Fig. 1F) licked for water reward according to the pairing relationship between the sample and test odors, which were separated by a delay period [as in ref. (*14, 15*), Fig. 1G and movie. S1]. In each trial, we delivered one of the two sample odors (S1, Ethyl propionate; S2, Propyl acetate), and following a delay period (3 or 6 sec), one of the two test odors (T1, Butyl formate; T2, Methyl butyrate). Mice were rewarded with water if they licked within a response window in the defined “paired” trials (S1-T1 or S2-T2), but not in the “unpaired” trials (S1-T2 or S2-T1, Fig. 1, G and H). Mice hardly licked during the delay period (fig. S1A).

We recorded brain-wide neuronal spiking with Neuropixels probes (*46*) (Fig. 1I, fig. S1, B and C) after mice were well-trained (criterion: performance of consecutive 40 trials > 75%; fig. S1D). Typically, two probes were simultaneously inserted in different regions of a mouse during each recording session (n = 223 probe insertions over 116 sessions in 40 mice, Fig. 1I). Neuronal spiking was detected and sorted into single units by Kilosort2 (*53*). A total of 33,028 single units (285 ± 13 units simultaneously) were successfully recorded from 62 brain regions (409 ± 92 units per region; 34 regions with more than 100 neurons; region as in the Allen Brain Atlas (*54*); fig. S1, C and E, full region-name list in table S1). Distances between inserted electrodes were larger than 1 mm to allow unambiguous identification of electrode tracks by histological reconstruction from fluorescently labelled probes (*46*) (fig. S1, C and E). Up to 12 electrode penetrations in 6 recording sessions were made in each mouse.

### Brain-wide activity waves encoding WM information

We found that many neurons could encode odor information during the delay period after the offset of external odor stimulus. These ‘memory neurons’ showed significantly different activity for sample odors during the delay period (an example neuron in Fig. 1, J and K; p < 0.05, Wilcoxon rank-sum test for selectivity throughout text). In a cross-validated support vector machine (SVM) decoding analysis, these neurons demonstrated significant decoding power for odor cues in the correct trials, which was decreased in the error trials (fig. S2A). The average firing rate of all neurons during the baseline period was 8.4 ± 0.1 Hz, whereas the firing rate of memory neurons in the preferred condition during the delay period was 10.5 ± 0.1 Hz.

The memory neurons are widely distributed across brain regions as shown in Figure 1L (24,667 neurons in 34 regions, only regions with more than 100 recorded neurons were included in the analysis). For a few brain regions that have been investigated previously with different recording techniques but similar behavioral-task design, such as AI (agranular insular area)(*15*) and PIR (piriform cortex)(*14*), the proportion of memory neurons are comparable across studies. Olfactory regions exhibited the highest proportion of memory neurons, such as TT (taenia tecta), AON (anterior olfactory nucleus), and PIR.

We then correlated the regional proportions of memory neurons with anatomical-connectivity results from the Allen Brain Atlas (*54*). As an illustration, the proportion of memory neurons in each region was positively correlated with the input strength from AON (Fig. 1M). Overall, the regional proportion of memory neurons showed a strong positive correlation with anatomical inputs from olfactory regions (e.g., AON, TT, and PIR, Fig. 1N), consistent with the causal importance of a sensory region in olfactory WM maintenance (*14*).

Most memory neurons exhibited significant odor selectivity only during some but not all time-bins of the delay period (fig. S2B; 26.8 - 27.5% for all delay duration), hereafter referred to as the “transient neurons”. We observed a small percentage of neurons (0.5 - 2.9%) exhibiting sustained selectivity across the 3-sec or 6-sec delay period following the sample offset, which were referred to as “sustained neurons” (fig. S2B, example in Fig. 1K). The transient neurons showed stronger correlation with behavioral performance than the sustained neurons, as shown by a larger reduction from the correct to the error trials in both normalized firing rate (Z-score, 0.23 to 0.09 for transient, 0.29 to 0.24 for sustained, fig. S2C) and Area Under Receiver Operating Characteristic curve (AUROC) (0.99 to 0.65 for transient neurons and 1.00 to 0.86 for sustained neurons, fig. S2D), consistent with previous results in the anterior agranular insular cortex (*15*). Therefore, brain-wide activity encoding olfactory WM information is dominated by transient patterns.

Although individual transient neurons only briefly encode information, population activity of these neurons could tile over the entire delay duration, readily revealed after sorted by <u>te</u>mporal <u>c</u>enter <u>o</u>f <u>m</u>odulation in firing rates (TCOM, Fig. 1O). Firing rate varied across trials, but we observed much higher correlation of TCOM for activity waves than for shuffled data (measured by variance explained or *r*^2^, in Fig. 1, O and P). Activity wave was significantly diminished during the inter-trial-interval (ITI) period when there was no need for memory maintenance (as shown by much lower variance explained in TCOM for held in vs. out in ITI period than that in delay period, Fig. 1P). Activity waves were correlated with behavioral performance, as shown by the differences in TCOM correlation between the correct and error trials (Fig. 1P). TCOM sorting for the WM-encoding neurons (27.8% of all neurons, 9,182/33,028) also revealed distinct activity waves following S1 and S2 odors (Fig. 1Q and R).

We next examined the correlation between wave timing and anatomical connections. The wave timing of a region was quantified by the averaged TCOM for the memory neurons within the region. Earlier wave timing for the memory neurons were correlated with more anatomical inputs from olfactory and hippocampal regions, therefore a negative correlation was observed [e.g., PIR, RHP (retrohippocampal region), Fig. 1S and fig. S3]. Thus, activity waves encoding odor information predominantly propagated from olfactory and hippocampus to other regions.

### Spike coupling underlying memory neurons and activity waves

To examine neuronal-activity dynamics supporting the second-scale activity waves, we examined millisecond-scale <u>s</u>pike <u>c</u>oupling (SC). Such SC represents sequential coactivation at a 10 millisecond-time scale of two neurons, which could reflect common inputs or be related to synaptic connections (*7, 55*). Previous studies have characterized task- and memory-related SC for neuronal pairs within local regions in WM tasks (*7, 56, 57*). However, the global dynamic pattern of cross-region SC in WM remains unclear.

Statistically significant SC was searched and detected between simultaneously recorded neurons using cross-correlation analysis (*7, 55*) (see Methods). The raster plot and spike cross-correlogram of an example SC neuronal pair (neuron 1 from taenia tecta, TT; neuron 2 from dorsal peduncular area, DP) are shown in Figure 2A and 2B. Many spikes of neuron 2 (blue bars) followed neuron-1 spikes (red bars) within 10 msec. Such <u>c</u>oupled <u>s</u>piking <u>p</u>airs (CSP) resulted in a statistically significant asymmetric peak with a short latency in the correlogram (p < 0.05, based on null hypothesis of independent Poisson distribution), indicating directional SC from the leading to the following neuron. Cross-region SC was inversely correlated with spatial distance between brain regions, which could be fitted with power law (Fig. 2C, see Method), reminiscent of the inverse correlation between anatomical connection and spatial distance (*58*). Within-region SC was also higher than cross-region SC (Fig 2, C and D).

**Fig 2.**
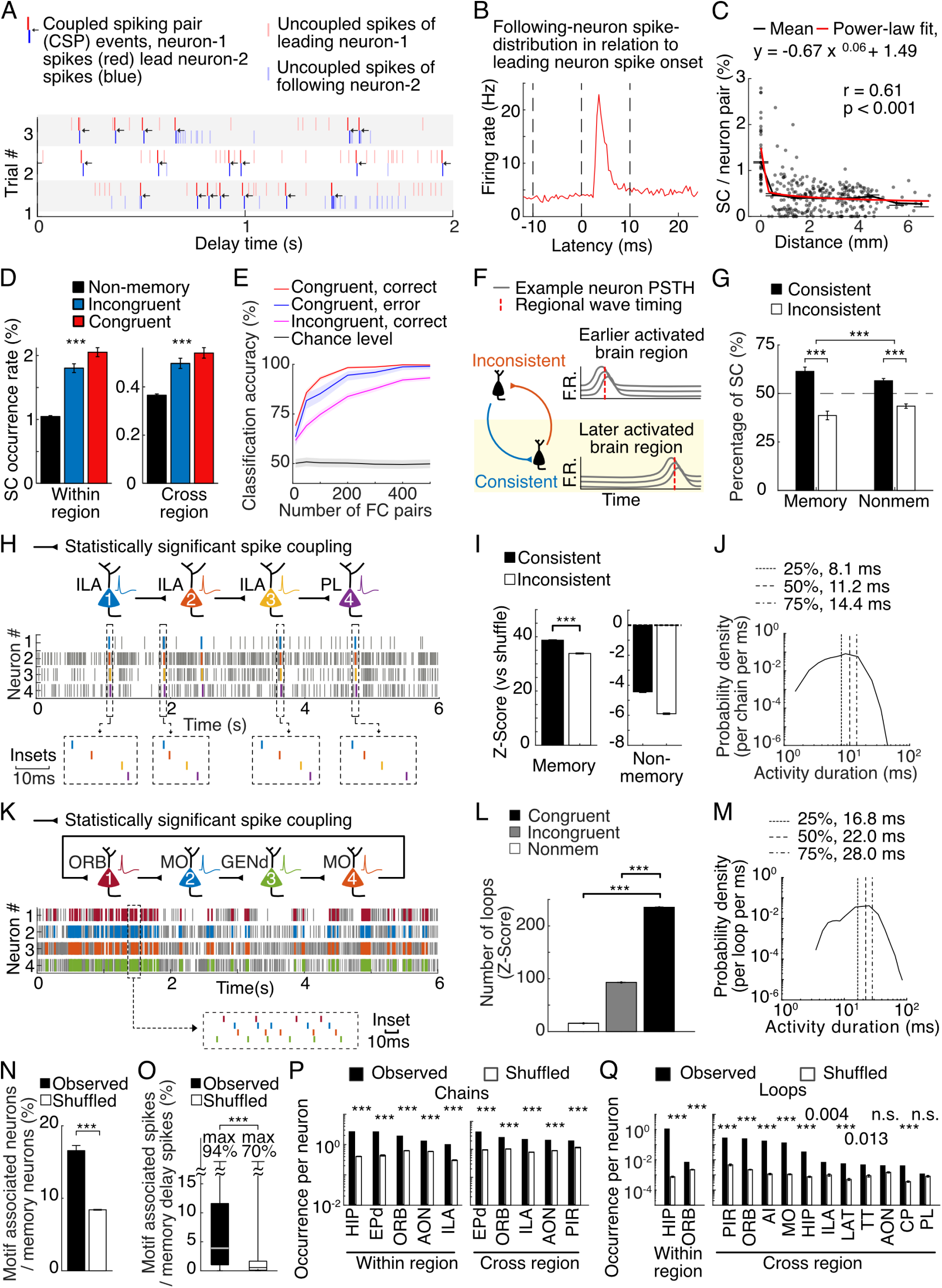
Memory-selective activity motifs of spike coupling, activity chains, and single loops. (**A**) An example pair of simultaneously recorded neurons with SC. Spike raster of three example trials from the two neurons with SC (red for the leading neuron; blue for the following neuron). Arrows indicate the coupled spiking pairs (CSP, with <10 msec latency). (**B**) Spike-correlogram showing the averaged spike frequency of the following neuron-2 in relation to spiking of the leading neuron-1. Vertical dashed lines indicate the period used to determine asymmetric peaks in spike-correlogram. (**C**) Relationship between SC rate and regional distance. (**D**) SC rate for the non-memory, incongruent-memory, and congruent-memory pairs, within and across brain regions. ***, p < 0.001 from rank-sum test. (**E**) Odor-sample classification accuracy in a cross-validated SVM decoder analysis, based on the CSP frequency of varied numbers of pairs of memory neurons, in the correct or error trials. X axis: number (10 to 500) of congruent or incongruent WM neuron pairs used. (**F**) Illustration of relationship between wave time and consistent or inconsistent SC. Cross-region SC was defined as ‘consistent’ if the leading neuron was in a region with earlier wave timing. (**G**) Proportion of consistent and inconsistent cross-region SC for memory and non-memory neurons. ***, p < 0.001, from Chi-Square tests. (**H**) Schematic drawing of an example activity chain. Above: Schematics of an example SC-connected activity chain with three neurons in ILA and one neuron in PL, with the direction indicated by the synapse symbols. Middle: Spike raster plot of delay-period activity during an example trial among the four memory neurons, with colored bars marking the spikes that constitute the activity chain, and gray bars indicating spikes that are not part of the activity chain. Below: Zoom-in view of several spiking sequences of the activity chain. (**I**) Shuffle-normalized ratio (Z-score) of observed consistent and inconsistent activity chains. ***, p < 0.001, rank-sum test across 1000 shuffle repeats. (**J**) Duration distribution for activity chains. The probability density is calculated by dividing the number of activity-sequences in each duration-time bin by the total number of chains and the bin width in milliseconds. (**K**) Similar to (H), but shows an example activity loop. (**L**) Normalized number of single loops of non-memory, incongruent-memory, or congruent-memory neurons, for either within- or cross-regional single loops. Z-score estimated from 1000 bootstrap resamplings of the shuffled dataset. Error bars, S.E.M. ***, p < 0.001, rank-sum test. (**M**) Duration distribution for single loops. (**N**) Proportion of memory neurons that participated in activity chains and single loops. ***, p < 0.001, Chi-Square test. (**O**) Proportion of spikes associated with activity chains and single loops relative to the total amount of spikes during the delay periods of selective neurons. ***, p < 0.001, rank-sum test (**P**) Fraction of chain-associated neurons among the total number of neurons in various regions, in observed and shuffled datasets, see figs. S6B and S6C for a complete list. ***, p < 0.001, probability from 100 shuffled repeats. (**Q**) Similar to (P), but for loops.

**Fig 3.**
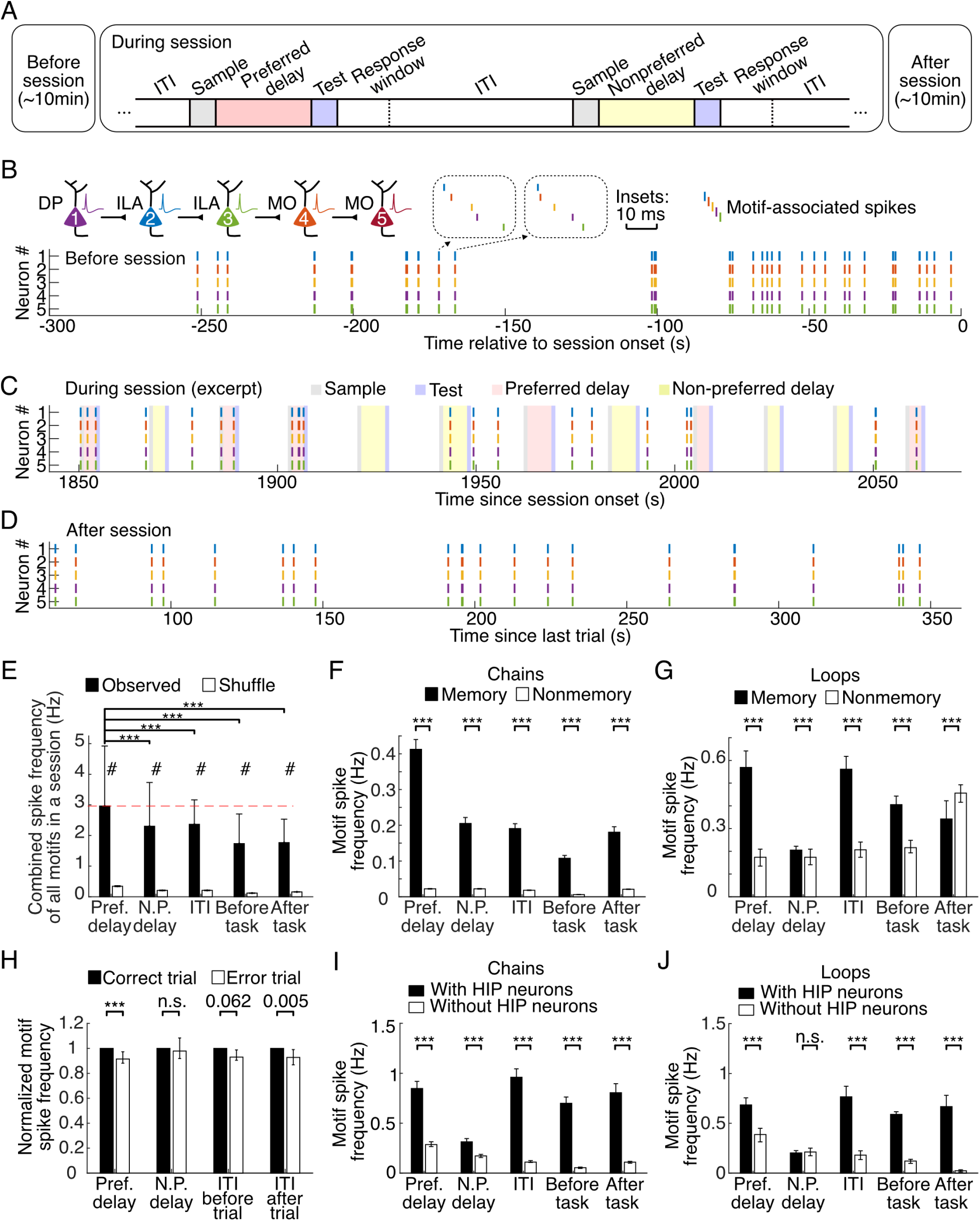
Replays of WM-associated activity chains and loops. (**A**) Schematic illustration of the 10-minute recording before the start (left) and after the end (right) of task execution, and the trial structure (middle) with 10-sec inter-trial intervals (ITIs) inside task execution. (**B**) An example of spontaneously replayed activity chain before the start of the task execution. (**C**) Occurrence of the same activity chain as in (B), during and between the delay periods. Color shades labeled above: corresponding task epochs. (**D**) as in (B), but after the end of task execution. (**E**) Combined spike frequency of all the activity motifs, during the preferred- and non-preferred delay, ITI, before- and after-task periods. Spike is counted only once if it participates in multiple loops or chains. Bars represent the across-session median, error bars represent the 95% confidence interval (C.I.) of the median from 1000 bootstrap resampling. Pref., preferred; N.P. non-preferred, throughout this figure. ***, p<0.001, signed-rank test; #, p < 0.001, from shuffle-normalized Z-score. (**F, G**) Spike frequency of activity chains (F) and single loops (G) composed of memory and non-memory neurons. ***, p<0.001, rank-sum test. Bars and error bars represent the median and 95% C.I. (**H**) Relationship between activity motifs and performance, during the delay period and off-delay period. ***, p<0.001, signed-rank test. Bars and error bars represent the median and 95% C.I. (**I**) Spike frequency from activity chains comprised of at least one HIP neuron, or chains without any hippocampal neurons during the delay or off-delay period. ***, p<0.001, rank-sum test. Bars and error bars represent the median and 95% C.I., as above. (**J**) As in (I), but for single loops.

SC analysis enabled us to examine the relationship between coupling strength and coding ability of the involved neurons. We tested it and observed stronger SC for the congruent neuronal pairs (Fig. 2D), defined as two memory neurons preferring to the same WM information (fig. S4A). The incongruent pairs, which involved two memory neurons preferring different memory information, exhibited lower SC than that between congruent memory neurons, but still higher SC than that between non-memory neurons (Fig. 2D).

To further reveal the behavioral relevance of CSP, we treated CSP as a computing unit and examined its coding ability. Odor-sample information could be decoded from CSP frequency by a cross-validated SVM analysis, better for the congruent than the incongruent pairs (Fig. 2E).

Although the spikes involved in congruent CSP only accounted for 19.9% of total spikes during the delay period for the involved neurons (8.6% - 41.5%, 25% - 75% range, fig. S4B), the contribution of coding ability of CSP in a memory-neuron pair was comparable to firing rates of a memory-encoding neuron (fig. S4C). The coding ability in the correct trials was also higher than that in the error trials (Fig. 2E). Therefore, sequential coactivation represented by CSP could maintain WM information in a manner correlated with behavioral performance.

We then examined whether the overall direction of cross-region SC in the memory-encoding neurons was consistent with that of activity waves. The wave timing for each region was quantified by the averaged TCOM of all the memory-encoding neurons in that region. If the regional wave timing of the leading neuron is ahead of that of the following neuron, we denoted this cross-region SC as ‘consistent’, otherwise ‘inconsistent’ (Fig. 2F). We found more consistent than inconsistent SC for the both the memory neurons and non-memory neurons, but the difference was significantly larger in the memory neurons (Fig. 2G). This result indicates that the overall direction of cross-region SC is in line with that of the activity waves.

### Activity chains and single loops linked by coupled spikes

Although we found more SCs consistent with regional wave timing, there were temporal gaps between the 1-10 msec SCs and the second-scale activity waves. We therefore examined whether SC could link neurons into activity motifs capable of more prolonged activation. Motivated by the synfire-chain theory (*27–29*) (Fig. 1C), we searched for activity chains connected by SC in the simultaneously recorded neurons. In the example shown in Figure 2H, three neurons from ILA (infralimbic area), and one neuron from PL (prelimbic area) were consecutively coupled (spike cross-correlograms of these neuronal pairs in fig. S5A). These four neurons were linked into an activity chain, as manifested by the sequential spiking of these neurons within 10-msec interval (Fig. 2H inset below).

For the cross-region activity chains of memory neurons, the direction could be either consistent with that of brain-wide activity waves (denoted as “consistent”), or opposite to that of waves (as “inconsistent”). We observed 3,044 consistent chains and 2,021 inconsistent chains, each comprised of 3 to 9 neurons. The consistent chains were significantly higher in occurrence than inconsistent ones (fig. S5B). The number of observed chains were much higher than the shuffled results, and the consistent chains are overrepresented to a greater extent than inconsistent chains (Z-score: 38.7 and 33.8 for consistent and inconsistent SC, respectively; Fig. 2I). Although we also observed non-memory chains, such occurrence was less than shuffled data (Z-scores −3.9 and −6.8 for the consistent and inconsistent non-memory chains, respectively, Fig. 2I). The continuous sequence in these chains typically lasted around 11.2 msec (median; 8.1 – 14.4 msec, 25% - 75% range, Fig. 2J). The within-region activity chains of the memory neurons were also much higher than the shuffled data and that of the non-memory neurons (fig. S5C).

Besides activity chains, another potential motif to support prolonged activation is recurrent loop (Fig. 1A), as suggested by the cell-assembly hypothesis (*21–23*). We therefore searched for the single loops composed of 3-5 neurons, which were linked back to an initiating neuron by SC. The diagram of an example single loop composed of four neurons in three regions is shown in Figure 2K (neuron 1→2→3→4→1). The zoom-in plot below Figure 2K clearly shows detailed sequential and looped spiking activity. The occurrence of single loops among the congruent-memory neurons was much higher than that among the incongruent and non-memory neurons (Z-score normalized to the shuffled chance level, Fig. 2L). The duration of a typical single loop was about 22.0 msec (median; 16.8 – 28.0 msec, 25% - 75% range, Fig. 2M).

The median spike frequency of the observed activity chains and single loops was 0.4 Hz (0.1 – 1.3 Hz, 25% - 75% range) throughout the delay duration (fig. S6A). We found that 16.6% memory neurons were involved in the activity chains and single loops (Fig. 2N). During the delay period, a median proportion (3.9%, median; 1.0% - 11.6%, 25% - 75% range) of all spikes is associated with chains and single loops among these memory neurons (Fig. 2O).

We then examined the regional differences in the memory neurons participating activity chains (measured by the number of neurons presented in activity chains divided by the total number of recorded neurons in each region). Some activity chains only involved the neurons of a single region, which were most prominent in the hippocampus, olfactory [EPd (endopiriform nucleus dorsal part), AON], and prefrontal [ORB (Orbital area), ILA] cortices (highest 5 regions in Fig. 2P left, full list in fig. S6B). Other activity chains involved cross-region SC, of which we observed higher occurrence of the neurons in the olfactory (EPd, AON, PIR) and prefrontal cortex (ORB, ILA, highest 5 regions in Fig. 2 right, full list in fig. S6C). The observed occurrence in these regions was much higher than the shuffled chance level.

For within-region single loops, the neurons of hippocampus and orbitofrontal cortex were highly involved (Fig. 2Q left). For the cross-region single loops (Fig. 2Q right), we found higher occurrence of the neurons in the prefrontal cortex (PL, ILA, ORB), anterior insular cortex (AI), olfactory regions (AON and PIR), motor cortex (somatomotor areas, MO), and hippocampus.

The observed occurrence was also much higher than the shuffled chance level.

### Replays of WM-associated activity chains and loops

We wondered whether the observed activity motifs could also occur outside of the delay period. We thus examined the activity motifs during the 10-sec inter-trial interval (ITI, Fig. 3A, middle) within task execution, as well as 10-minute recording epochs before the start and after the end of task execution (Figure 3A, left and right).

Intriguingly, many motif activities could be observed during these periods outside of delay period (termed ‘off-delay period’). In the example recording shown in Figure 3B-D, the activity chain was composed of five neurons from DP (dorsal peduncular area), ILA, and MO regions. Even before the task excitation, we already observed occurrence of this activity chain (Figure 3B). During task execution, this activity chain occurred more often during the delay period following the preferred sample (in 4 out of 6 preferred sample trials, comparing to 1 out of 6 non-preferred sample trials, pink vs. yellow background, Fig. 3C). Importantly, this activity chain also occurred during the ITI (in 6 out of 13 ITI, Fig. 3C). After task execution, we still observed occurrence of this activity chain (Fig. 3D). The overall frequency of this activity chain before task execution, during delay period, ITI, and after task execution was 0.7 Hz, 1.9 Hz, 0.4 Hz and 0.4 Hz, respectively.

We then computed the replayed frequency for all the recorded activity motifs. The observed motif occurrence significantly exceeded the shuffled control in all behavioral epochs (observed vs. shuffle in Figure 3E), thus replay significantly occurred outside of the delay period. The frequency of the replayed activity motifs was about 2/3 of that during the delay period, as shown by the combined frequency of spikes belong to any memory-associated activity motifs during either delay or off-delay period (cross-session statistics in Fig. 3E). The replayed activity motifs were highly related with WM maintenance, as shown by the large differences between chains/loops composed of memory and non-memory neurons (cross-motif statistics in Fig. 3, F and G). Notably, replay frequency of the activity chains during ITI was lower than the occurrence frequency during the delay period following preferred sample (termed as ‘preferred delay’, Fig. 3F), but replay of the single loops during ITI was actually similar to that of preferred delay (Fig. 3G).

We also examined whether the occurrence of activity motifs was correlated with behavioral performance. Indeed, during the delay period, we observed slightly weaker activity motifs in the error trials than in the correct trials (Fig. 3H, delay period). We also observed slightly weaker replayed activity motifs during ITI of the error trials (Fig. 3H, ITI). During the ITI, the replayed motif spikes were not associated with sample odor of the immediate previous trial, nor with the next following trial (fig. S7A). Such lack of activity difference during ITI is consistent with a small behavioral effect of previous trial history, as estimated by a general linear model (fig. S7B).

Because of the importance of hippocampal neurons in activity motifs during the delay period (Figure 2P and Q) and in replay during either sleep and awake (*32, 33, 35, 37, 38*), we further examined the involvement of hippocampal neurons in replay of activity motifs. Comparing to motifs comprised of at least one HIP (hippocampal region) neuron, activity from motifs without any hippocampal neurons indeed occurred in significantly lower frequency for both activity chains and single loops during the delay or the off-delay period (Fig. 3, I and J). Intriguingly, replayed activity chains and single loops involving hippocampus occurred at a similar level comparing to that during the delay period following preferred sample odor (termed ‘preferred delay’, Fig. 3, I and J). This was in sharp contrast to the results for all motifs (Fig. 3, F and G). Moreover, during the delay period following non-preferred sample, activity motifs involving hippocampus occurred at a lower frequency (Fig. 3, I and J). Therefore, hippocampal neurons are highly involved in the activity replay of spiking motifs.

### Hierarchical organization of activity chain, single loops, and nested loops

We sought out to search whether the activity chains and single loops could be further linked into more complex structures, an example of which was the nested loop shown in Donald Hebb’s drawing of a hypothetical assembly (*21*) (Fig. 1A). In such nested loops, the activity of different loops could be connected by commonly shared neurons. Figure 4A shows an example nested loop that we observed, composed of 6 neurons from 4 regions. Black synapse symbol indicates directional SC, whereas colored arrows indicate observed occurrence of single loops. Sequential activity of an example trial for this nested loop is shown in Figure 4B. Another example nested loop is shown in Figure 4C, with 18 neurons of congruent WM selectivity linked into 28 chains and 7 loops, which were all integrated inside a nested structure. Snapshots of activity loops in this example nested loop are shown in Figure 4D. The raster plot of spiking of chains and single loops involved in the nested loop (fig. S8A) showed sequential alternative activation of various motifs at a 10-msec resolution. The spike frequency associated with this nested loop was approximately 40Hz (fig. S8A).

**Fig 4.**
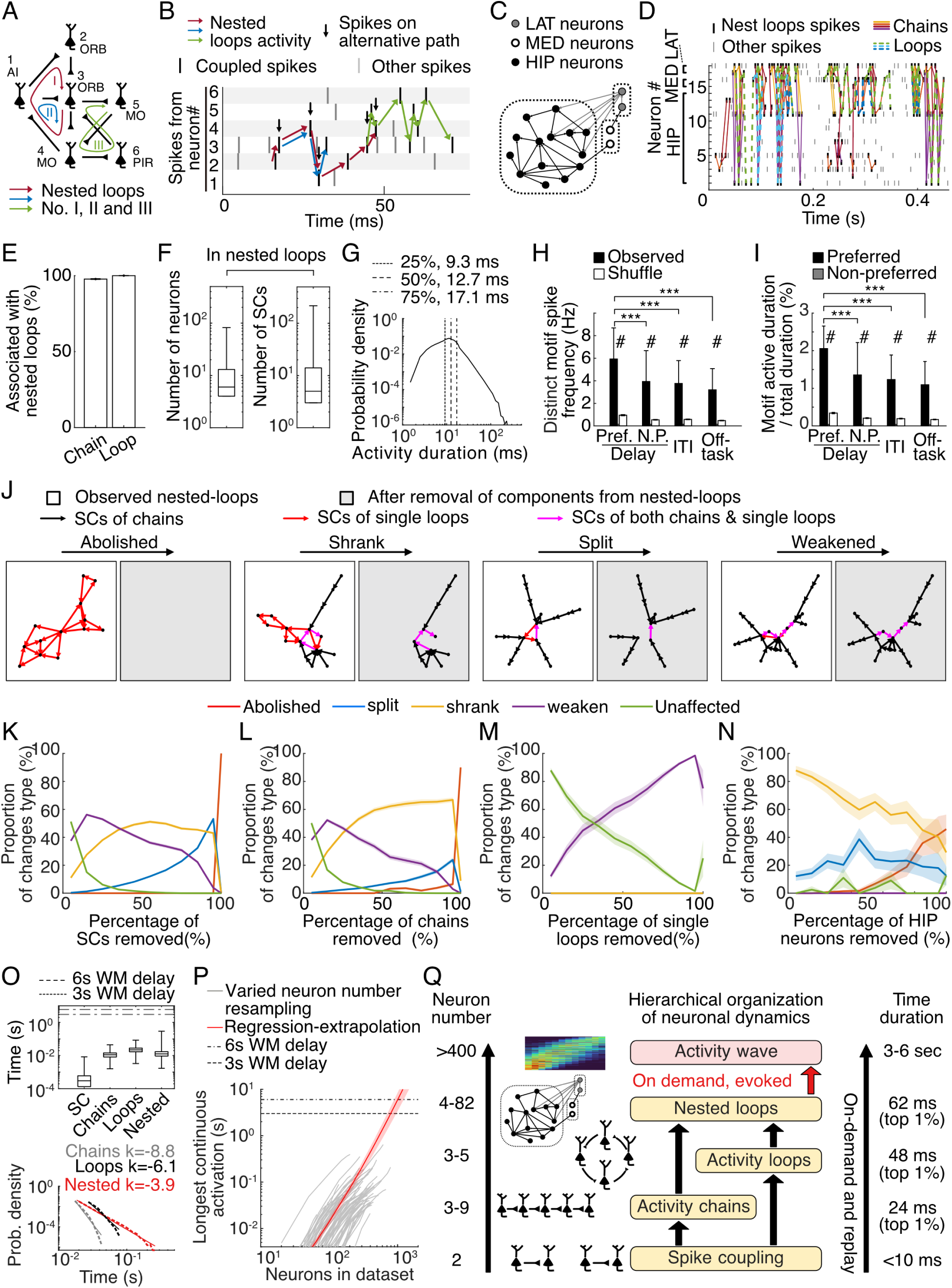
Nested loops and hierarchical organization of cross-region sequential spiking. (**A**) An example nested loop that is similar to the example assembly proposed by Donald Hebb (*21*). Neuron ID and region name were listed. Synapse symbols indicate SC, and colored lines indicate three example alternative activity loops. (**B**) Spike raster of an example trial of alternative loop activity for the example nested loop in (A), with the same color-coded arrows indicating spiking sequence within respective loops. (**C**) Another example nested loop, comprised of 18 memory neurons, 28 chains, and 7 single loops. (**D**) Raster showing example chained-loops activity for (C). (**E**) Proportion of activity chains and single loops that belonged to nested loops. (**F**) Statistics of the number of neurons (nodes in graph theory) and the number of SCs (edges) in the nested loops. (**G**) Distribution for the duration of consecutive sequential spiking in the nested loops. (**H**) Spike frequency associated nested loops during the WM delay, ITI, and off-task (before and after task execution) periods. Pref., preferred, N.P., non-preferred, throughout this figure. Bars and error bars represent median and 95% C.I., as above. ***, p < 0.001, signed-rank tests. #, p < 0.001, from shuffle-normalized Z-score. (**I**) Proportion of time that were covered by continuous nested loops activity during WM delay, ITI, and out-of-task periods. Bars and error bars represent median and 95% C.I., ***, p < 0.001, #, p < 0.001, as above. (**J**) Illustration of four potential changes to nested loops after element removal. (**K-L**) Changes after gradual removal of increasing number of SCs (K), activity chains (L), single loops (M), and neurons located in the HIP region (N). (**O**) Above: An overview of the time-constant distribution of activity motifs in various forms. Whiskers, full range. Below: Fit the tails of the duration distribution (above) to power-law functions. Dashed curves represent observation, solid lines represent fit prediction. Numbers above: exponent parameter from curve fitting. (**P**) Grey curves depict the relationship between systematically varied numbers of neurons and the average longest activation-duration across multiple sessions. Red curve represents the linear regression and extrapolation of this relationship, shadow indicates S.E.M. (**Q**) Hierarchical organization and replay of neuronal dynamics underlying memory-specific brain-wide activity waves and working-memory maintenance.

We then searched for such nested loops in all the recorded neurons. We found nested loops in 76.9% of sessions with at least one motif. Most of the activity chains and single loops detected in Figure 2 were associated with some nested loops (> 97.7% for both types of motifs, Fig. 4E). The nested loops were composed of 4 to 82 neurons, which were connected by 3 to 217 SCs (Fig. 4F). From a graph-theory point of view (*59*), the connected density within a nested loop was 16.7% (median, as defined by the number of actual SC divided by that of all possible SC; 9.9% - 25.0%, 25% - 75% range; fig. S8B). The duration of spiking occurrence in the nested loops typically lasted 12.7 msec (9.3 – 17.1 msec, 25% - 75% range, Fig. 4G). If a longer delay was allowed in connecting activity chains and single loops, increased duration of consecutive activation was observed in the nested loops (Fig. S9A). Among these nested loops, the neurons in the endopiriform, hippocampus and orbitofrontal cortex exhibited a higher level of connectivity, as quantified by the average node degrees (fig. S9B).

Such nested motifs were observed much more frequently than the shuffle level, exhibited memory selectivity during the delay period, and were replayed outside the delay period (Fig. 4, H and I, example in Fig. S10). The spike frequency associated with the replayed nested loops was lower than that during the delay period (Fig. 4H). The duration covered by the replayed nested loops was also significantly shorter than that during the delay period (Fig. 4I). Both phenomena were consistent with the reduced frequency of the activity chains and single loops during the off-delay period (Fig. 3, F and G, with the exception of single loops during ITI).

To further dissect the contribution of different elemental motifs in the integrity of the nested loops, we performed the analysis to gradually remove specific components and then examine the changes to the nested loops. Several types of changes were observed after removing different proportions of SCs. First, a nested loop could be entirely abolished, without any neurons connected by SC (‘Abolished’ in Fig. 4J). Second, a nested loop could be reduced in size, yet still connected into one loop (‘Shrank’ in Fig. 4J). Third, one nested loop could be split into two or more smaller loops (‘Split’ in Fig. 4J). Finally, a nested loop could be weakened in the number of connected edges, yet no change to the number of participating neurons (‘Weaken’ in Fig. 4J). After removal of increasing number of SCs, a nested loop was initially weakened in the number of edges, then gradually shrank in the node size, then further split, eventually abolished (Fig. 4K). Removing activity chains exhibited similar patterns (Fig. 4L). Removing single loops, however, predominantly induced weakening in the edges (Fig. 4M). Interestingly, removing hippocampal neurons exhibited entirely different patterns, with little unaffected nested loops and a large proportion of shrank. Removing hippocampal neurons also induced split and abolished nested loops faster than other manipulations (Fig. 4N). Thus, SC and activity chains, single loops, as well as hippocampal neurons contributed differently to the formation of nested loops.

There was a progressive increase in the long end of the time-constant distribution for the observed activity motifs (24.4 msec, 47.8 msec, 61.8 msec for the longest 1% of activity chains, single loops, and nested loops, respectively, Fig. 4O, above). The falling tail in these distributions could be fitted to a power-law function, P.D.(*t*) = α*t*^k^, where the probability density of a motif persisting for *t* second is proportional to that to the power of k. Increasing k values for activity chains, single loops, and nested loops were consistent with the notion of climbing levels along a hierarchy (Fig. 4O, below). Given that only hundreds of neurons were simultaneously sampled from 70 million neurons of the mouse brain, we suspected that the overall duration of such continuous stream of connected motifs should be drastically increased with a higher number of simultaneously recorded neurons. Systematic removal of neurons indeed revealed a linear relationship between the longest activation duration and the number of simultaneously recorded neurons (Fig. 4P). Extrapolation curve showed that spiking sequences composed of these activity motifs from thousands of neurons could outlast the entire delay length (3 or 6 sec).

## Discussion

Taken together, our Neuropixels recordings revealed a hierarchical organization of activity motifs, composed of sequential spiking from the neurons both within region and across multiple brain regions. From bottom to top, the motifs of SC, activity chains, single loops, and nested loops were hierarchically organized, with progressively increasing time constant and the number of participating neurons (Fig. 4, O and Q). Such hierarchically organized activity motifs were replayed before and after task execution, as well as during ITI. Upon external application of olfactory cue and on demand for WM maintenance, activity motifs were either increased or decreased according to their memory selectivity, which could support memory-associated activity waves and mediate perpetual working memory.

Behavior-related activity patterns could be replayed in sleep or awake state, prominently in the hippocampus and neocortex, largely accompanied with slow-wave ripples and with time compression of past experience (*32–41*). However, it remains unclear whether and to what extent the working memory-related spiking can be replayed beyond the delay period. To our surprise, we observed replay of cross-region activity motifs, before and after the task execution, as well as during the inter-trial interval. Moreover, stronger activity loops and single loops were replayed among the neurons encoding the same information, for most of the time of off-delay periods (except for the single loops after task execution) (Figure 3F, 3G). Notably, single loops occurred at the similar frequency during ITI as in the preferred delay period (Figure 3G). Furthermore, slightly more activity replays were observed during the ITI of the correct trials versus the error trials (Figure 3H). Learning could shape the neural circuits into replayed cell assemblies following ‘fire-together wire-together’ principle (*21*). The involved plasticity rules (*60*) and the changing nature of activity motifs during learning remain to be determined. The repetitive occurrence of activity motifs during learning of the WM task might lead to specific strengthening of motif-related synaptic contacts, which further lead to spontaneous emergence of such activity motifs under the head-fixed behavioral context, either before or after task execution, as well as during ITI. Such activity replay could facilitate the reactivation and consolidation of task rule-related memories (*39, 42, 45*), and constructing internal representations to improve task performance (*44, 45*).

Hippocampus is essential for acquisition of episodic and spatial memory (*61*), and is critical for cognitive map (*32–41*). Although best known for the deficits in long-term memory (*61*), hippocampus lesion also induced WM deficit under more challenging situations, for instance with more items in WM or a long delay period (*62*). Animal studies showed that optogenetic suppression hippocampus-to-mPFC (*63*) and entorhinal-to-hippocampus projections (*64*) impaired spatial WM performance, and hippocampal neurons exhibited transient patterns of activity in encoding olfactory and temporal information during the delay period (*16*). Our results demonstrated that hippocampal neurons played important roles in the integrity and replay activity motifs (Figure 2P, 2Q, 3I, 3J, 4N). Moreover, the hippocampus-associated chains and loops showed similar occurrence frequency as that during preferred delay, and showed suppressed occurrence during non-preferred delay (Fig. 3, I and J). Thus, replay of activity motifs involving hippocampus is prominent even outside of delay period, and is actively suppressed following the non-preferred sample odor. Such replay might be important in organizing global brain activity in episodic memory and cognitive map. The cell-type and structural foundation of these contribution of hippocampal neurons remain to be determined in future studies.

We noted that the replayed single loops after task execution for non-memory neurons actually occurred more frequently than that for memory neurons (Figure 3G), whereas in all other conditions we observed a higher frequency of replayed motifs for memory neurons. The function of this increased non-memory loops after task execution remains to be determined, which might be related with anticipation of returning to home cage or after-task water supplement (see Methods).

Cross-region SC in our study was based on spike correlogram analysis on extracellular-recording results. Although SC was likely to be related with synaptic connection (*7, 55*), more direct techniques are necessary to characterize the properties and short-term plasticity of mono-synaptic connections. Future investigations with such techniques, for example combined whole-cell and extracellular recordings *in vivo* (*65*), are required to reveal the mono-synaptic basis of global cell-assembly dynamics underlying WM.

The activity of transiently selective neurons was tiling over the delay period (Fig. 1). Such transient delay-period activity has also been reported in insular cortex and hippocampus for olfactory WM tasks (*15, 16*). However, attractor network dynamics through sustained firing in individual neurons were found to play important roles in other WM tasks, e.g., in motor-planning oriented task (*18*). In our recordings, we observed ‘wave-and-stay’ phenomenon (about 5% memory neurons shown on top of Figure 1Q and 1R), consistent with the ‘ramp-and-stay’ phenomenon observed in delayed-response task in ALM neurons (*5, 10, 18, 19*). Thus, neuronal activity patterns underlying WM might be flexible for sensory modality, task demand, and training history.

In summary, spiking motifs are hierarchically organized and replayed, and are modulated on demand to support activity waves and perceptual working memory.

## Acknowledgments

We thank Drs. Muming Poo, Liping Wang, Bin Min, Yunzhe Liu, Ninglong Xu, Tianming Yang, Min Xu, Chun Xu and Yong Gu for critical comments on the project and manuscript. We thank the Center for Data and Computing in Brain Science, light-microscopy and electron-microscopy imaging core facilities of CEBSIT/ION in obtaining corresponding results.

## Funding

Innovations of Science and Technology 2030 from the Ministry of Science and Technology of China 2021ZD0203601 (CY)

National Key R&D Program of China 2019YFA0709504 (CY)

National Natural Science Foundation of China 31827803 and 32161133024 (CY)

Strategic Priority Research Program of the Chinese Academy of Sciences XDA27010400 and XDB32010100 (CY)

Shanghai Municipal Science and Technology Major Project 2018SHZDZX05 and 2021SHZDZX (CY, XZ)

Lingang Laboratory LG202105-01-01 (CY, XZ)

Shanghai Pilot Program for Basic Research-Chinese Academy of Science, Shanghai Branch JCYJ-SHFY-2022-010 (CY)

## Author contributions

E.H., D.X. H.Z. performed behavioral and recording experiments, analyzed results, and plotted figures; Z.C. performed recording and other experiments; C.Y. helped in data analysis; Y. C., R.H, H.F. and T.C. helped in various experiments; X.Z. and C.L. conceived the project, supervised experiments, analyzed results and wrote manuscript.

## Data and materials availability

All data and code used in the analysis have been deposited publicly and can be accessed at https://datadryad.org/stash/share/Z7wUPpxCmJWM621LgU4TjNxakIlV_CFl-op1nc4n9lo and https://gitee.com/XiaoxingZhang/pixels.

## Materials and Methods

### Experimental model and subject details

Some of the following methods are similar to those previously published (*11, 14, 66, 67*). C57BL/6J Slac male adult mice were provided by the Shanghai Laboratory Animal Center (SLAC), CAS, Shanghai, China. All mice were healthy, male, 8–12 weeks in age, and 20–30 g in weight at the start of the training. Mice were group-housed (4-6 per cage) under a 12-hour light-dark cycle (light on from 6:30 a.m. to 6:30 p.m.). The electrophysiological data presented here were gathered from 40 mice in order to obtain at least 30 brain regions with more than 100 recorded neurons. All experiments were performed in compliance with typical animal care standards and have been approved by the Institutional Animal Care and Use Committee of the Institute of Neuroscience, Chinese Academy of Sciences (Shanghai, China, Reference Number NA-014-2019).

### Behavioral setups

We utilized the delay-varying olfaction delayed paired association (ODPA) task in this study as previously reported (*11, 14, 15, 66*). In brief, the olfactometry apparatus was custom-built around a PIC Digital Signal Controller (dsPIC30F6010A, Microchip, Chandler, AZ), and was used to control olfactory cues and water delivery by switching solenoid valves and to detect lick responses with an infra-red beam break detector. The odor-delivery assembly was 3D printed and included an air-and-odorant-mixture nozzle, a lick port, and a beam breaker assembly. The assemblies were placed in front of the mice during the training and recording. The air flow rate was controlled at a constant rate of 1.35 L/min. Liquids of butyl formate (Cat. Num. 261521, from Sigma-Aldrich for all odors), methyl butyrate (246093), ethyl propionate (112305), and propyl acetate (133108) were distributed in air-tight bottles to produce odorant vapor. Polytetrafluoroethylene tubes were used to link the bottles to solenoid valves. During odor delivery, individually controlled air flow passed through the bottle with the odorant vapor, which was then mixed with air at a 1:10 concentration (v/v). The concentration of odors was measured using a photoionization detector (200B miniPID, Aurora Scientific Inc., St., Aurora, Canada), and the concentration during the delay fell to the baseline level within 1 sec after valve shut-off (*11, 14, 15, 66*). Behavior events and timings were recorded by a desktop computer using in-house developed software (https://github.com/zhangxiaoxing/SerialJ).

### Behavioral training

In the ODPA task, one of the two sample odors (S1: Ethyl propionate or S2: Propyl acetate) was presented for 1 sec at the start of a trial, followed by a delay duration of 3 sec or 6 sec. Mini-blocks of four trials were switched between each delay duration. After the delay period, one of the two test odors (T1: Butyl formate or T2: Methyl butyrate) was presented for 1s. After the end of the test odor offset, mice could respond in the response window lasting for 1s. The inter-trial interval was 10s. One sample odor was paired with a particular test odor (S1-T1; S2-T2). Mice were trained to lick in the response window only in paired trials. Hit and false alarm were defined by licking in the response window of the paired and unpaired (S1-T2; S2-T1) trials, respectively. Miss and correct rejection were defined as the absence of licking in the response window of the paired or unpaired trials, respectively. A reward of 2.5 µL of water was triggered immediately after the hit responses. Mice were neither rewarded nor punished in the false alarm and missed trials. There were 60 training blocks every day and each block contained 4 trials, interleaved between 3 sec and 6 sec delay duration. In total, mice performed 240 trials in each recording session. In the majority of the data analysis, only trials with correct responses within well-trained performance windows (above 75% performance rate for 40 consecutive trials) were included; however, in the correct-error trial comparisons, error trials within or outside the well-trained window were included to compensate for the smaller number. After starting behavior training, mice could obtain water as the reward for performing the task. After each training session, more water was given to mice whose body weight was less than 80% of their baseline weight.

Mice were trained in habituation and shaping sessions before the ODPA task. In the habituation session, mice were head-fixed in a tube and learned to obtain water by licking the water port. Water (2.5 µL) was delivered immediately after each lick, with a minimal interval for water delivery of 200 msec. The habituation sessions lasted for 2 days. In the shaping sessions, only the paired condition was presented, and mice could obtain water in all trials as long as they licked the water port in the response window. Shaping could last 2-5 sessions. The full training of ODPA started after reaching 75% performance in the shaping sessions.

### Surgery and Stereotaxic implantation of Neuropixels probes

Before the start of surgery, mice were anesthetized with sodium pentobarbital (0.1mL/g body weight) *via* intraperitoneal injection and placed in a stereotaxic instrument. All surgical tools and materials were sterilized by autoclaving or ultraviolet radiation for more than 20 min. During the surgery, a heating blanket was used to keep the anesthetized mice warm. Scalp, periosteum and other associated soft tissues were removed, and a custom-designed stainless-steel plate (fig. S1x) was fixed onto the posterior part of mouse skull with tissue adhesive (1469 SB, 3M, Maplewood, MN) and dental cement. For precise insertion of Neuropixels probes, the designed entry sites were marked on the skull. Then, a thin layer of dental cement was applied on the skull for better protection of skull surface. After the surgery, mice were kept on the heating pad until they woke up and the antibiotics (0.2mL 3.125mg/mL ceftriaxone sodium) was intraperitoneally injected for consecutive 3 days. At least 7 days of recovery was allowed before starting behavioral training.

On the day before recording, the skull under marked sites was removed with a cranial drill to make cranial windows (0.5-1.5mm in diameter), without removing the dura matter. For brain protection across days of recording, the cranial windows were covered first in 1.5% agarose, then silicone elastomer, and lastly dental cement. Mice were anesthetized with isoflurane in this process.

On the recording day, dental cement, silicone elastomer, and 1.5% agarose were removed from the skull in awake mice and the Neuropixels probe was inserted after the removal of dura mater in each cranial window. To label the probe tracks, each probe was dyed repeatedly with DiI, DiO and DiD (Invitrogen V22889, Thermo Fisher Scientific, MA) to label it as red, yellow and blue, respectively. When the tips of the probes touched brain surface, the depth of the stereotaxic instrument was set to zero. At the same time, the electrophysiological signals from the leading channels on the probes changed significantly in most cases. The probes were then gradually lowered into the brain until it reached the pre-defined target depth as measured by the stereotaxic instrument. Then we waited for thirty minutes before recording, to recover the brain from probe insertion and minimize probe drift during recording. The first 10 min of recording were used to record mice’s baseline activity before the start of behavioral training. A camera was used to capture mouse facial movements during the training.

After the task, the recording continued for an additional 10 min. After the recording, the probes were gradually pulled out of the brain. Then, 1.5% agarose, silicone elastomer, and dental cement were applied to the skull again, before releasing mice to home cage. The probes were cleaned with filtered pure water (>18 MOhm) and isopropyl alcohol after each insertion.

### Electrophysiology recording and spike sorting

Electrophysiology recording was carried out with the typically two Neuropixels probes simultaneously and the SpikeGLX software (https://github.com/billkarsh/SpikeGLX). The reference and ground pads on each probe were connected with soldered wire and all the ground wires were further connected together at a single point on a common ground connector block, which was further connected to the skull through the metal headplate. The action potential-recording gain was set to 1000, the local field-potential gain was set to 50, and the references in SpikeGLX software were set to the tip of each probe. The behavior controller (see Behavioral setups) outputs odor onset/offset and lick detection TTL signal in real-time on 3 BNC connectors. A custom-made embedded controller based on the PIC12F1572 MCU (Microchip, Chandler, AZ; code available at https://github.com/zhangxiaoxing/12F1572SyncEncoder) sampled the task and behavior signals at 1MHz to produce serialized synchronize signal in 4-bit words at 500Hz. The serialized signal was then connected to the SpikeGLX application through the built-in SMA port on the Neuropixels base-station. This serialized signal was synchronized to the electrophysiology data acquisition clock and were recorded in the same data file as an auxiliary track.

Spikes were sorted offline based on the waveform patterns. Single units were identified automatically using Kilosort2 (*53, 68*). Units with a contamination rate larger than 10% or firing rate lower than 1Hz were discarded.

### Analyzing samples-odor selectivity

Raster of spikes times were converted into firing rates (FR) of 1 second time bins that aligned to the sample-odor onset. The FR in the correct trials during the well-trained phase were then categorized into groups according to the sample odors and delay duration. Then, two Wilcoxon rank-sum tests were conducted for the time bins during the WM-delay following sample odor offset: (1) between one sample-duration group (i.e., S1 odor and 3s duration) and the rest 3 groups mingled (1 vs. 3) for each class, (2) between all the trials using Ethyl-propionate (S1) as sample odor and all the trials using Propyl-acetate (S2) as sample odor. If the FR within any of the 1-sec bins was significantly different in the 2 tests, the neuron was deemed memory-selective or memory neuron. The preferred task condition is determined by the highest averaged FR among the 4 conditions. Neurons that show no significant difference in both tests were identified as non-memory. The neurons showing significant selectivity in 1 to 2 bins in the 3-sec delay trials and 1 to 5bins in the 6-sec trials were characterized as transient, whereas the neurons showing significant selectivity in all 3 or 6 bins respectively were characterized as the sustained.

### Activity SVM Classifier analysis for firing rates in memory neurons

The SVM classifier analysis for the firing rates of memory selective neurons were carried out in MATLAB with the *fitcsvm* function. In each repeat of cross-validation, firstly, 50 (for fig. S2 A, C and D) or 5 to 500 (for fig.S4C) memory neurons were randomly chosen from the dataset. Then, 20 correct trials and 2 error trials for each label (odor S1 or S2) were randomly selected for each neuron in the corresponding sessions. The firing rates of the neurons in the selected trials were normalized using the min-max (range) method and 10-fold classification cross-validation (CV) was performed across the correct trials. Within each CV repeat, the correct-trials-trained model also classified corresponding firing rates in the error trials. The CV process was further repeated 25 times to obtain 1000 individual classification results (4 classification × 10 folds × 25 repeats) in both correct and error trials.

### Histology

After finishing the experiments, mice were deeply anesthetized with sodium pentobarbital (120mg/kg) and then perfused transcardially with 20 mL 0.9% NaCl solution followed by 20 mL paraformaldehyde (PFA, 4%, w/v) solution in phosphate-buffered saline (PBS). The brains were fixed in PFA solution for one or two nights and then transferred to PBS at 4℃ for storage. Coronal slices were obtained at 80-100 μm in thickness with a vibratome (Leica) and were incubated with DAPI (C1002, 1:1000, Beyotime, Shanghai, China) for 15 minutes then rinsed with PBS. Then slices were mounted and covered with coverslips. Slices were imaged at 10x magnification individually with an Olympus VS-120 fluorescence microscope. The color channels used for imaging were matched with dyes in each brain (DiD: 645nm; DiI: 535nm; DiO: 488nm; DAPI: 365nm).

### Histological probe localization

The pipeline of probe track localization was adapted from ref. (*47, 69*). Briefly, raw images were down sampled to 1/5 of their size. For each channel, brightness and contrast were adjusted manually to better visualize probe dye and brain areas. Then a custom-developed macro-script (NPpreprocess.txt, also see the code availability section) for FIJI (http://fiji.sc/) was used to preprocess and transform the images into TIFF format, which were further registered to Allen Institute Common Coordinate Framework (Allen CCF v3, http://download.alleninstitute.org/informatics-archive/current-release/mouse_ccf/). Custom-developed codes were used for further analysis (https://github.com/StringXD/SmartTrack-Histology). Individual slice images was resized to match the CCF template, then paired landmarks (∼10 points) in both the histology slice and the CCF template were labeled manually so that slice images could be registered to the CCF template. Since the probe tracks could span across multiple slices, after labeling the probe landmarks manually, a straight line was fitted with these points and served as the probe track. The electrophysiology data were associated with the corresponding histology tracks based on the mouse ID, the dye of the track, probe insert location and angle. Along with the tip location, electrophysiological landmarks in multi-unit spike raster, single unit spatial density and peristimulus-time histogram were included for positioning the recoding sites on a track-by-track basis. The Allen CCF anatomical localization along each track was exported to a list for further analysis.

### Temporal center of modulation and wave timing

The temporal center of modulation is defined in memory-selective neurons as follows. For each neuron, the delay-period firing rates in the preferred or nonpreferred task trials were separately averaged, labeled as FRP and FRN. The base-level is defined as the mean of FRP and FRN. Then, firing rates in the 3-sec or 6-sec delay period were averaged in 250 msec time bins across all correct preferred task trials. The base-level is then subtracted from the values to produce the modulation vector *w*. The temporal vector *t* is the time of the later boundary for each time bin. The temporal center of modulation (TCOM), in units of seconds, is then calculated as the sum of *w*_bin *_ t_bin_, divided by the sum of *w*_bin_, for bins 1 to 12 (in 3 sec-delay trials), or 1 to 24 (in 6 sec-delay trials).

### Brain-regional projection density and correlation to memory neurons

A custom developed script interfacing with the Allen Software Development Kit (https://allensdk.readthedocs.io/) was used to retrieve the structure-level projection data from the Allen Mouse Brain Connectivity Atlas. For each of the available grey matter injection sites (source brain regions) in both hemispheres, the *projection_density* data corresponding to each target brain region in a list of experiments is stored in a three-dimensional (inject-site vs. target-region vs. experiment-id) matrix. In the subsequent analysis, the median along the experiment-id axis is taken as the representative projection density.

For the correlation between projection density and regional proportions of memory-selective neurons (Fig. 1H and G, fig. S3), the distribution of projection density from a source region, such as the AON, to multiple target regions was compared to the proportions of memory-selective neurons in these target regions. Pearson’s correlation on log10-mapped projection density and memory neuron percentage data yields the correlation coefficient r. (Fig. 1H and fig. S2). This procedure is performed on additional available source brain regions to generate the brain-wide distribution data. Similar procedures were followed for the correlation between projection density and regional wave timing, with the distinction that the regionally averaged wave timing data were not log-transformed.

### Data replicates and statistical tests

In the present study, statistics on observed data and associated graphical representations are derived from biological replicates (e.g., Fig. 2D). The shuffled controls were based on technical replicates and were clearly labeled when used (e.g., Fig. 2P). In the resampled comparison of observations (e.g., Fig. 1P), both biological replicates (neurons) and technical replicates (resampling) were incorporated.

For the statistical tests between real-numbered data values, Wilcoxon signed-rank test was used to compare repeated measurements of the same group of subjects across multiple conditions (e.g., Fig. 3E, observed). If the subjects were biological replicates and not matched, the Wilcoxon rank-sum test is applied (e.g., Fig. 3F). For the comparison between observed and shuffled data (e.g., Fig. 2P), a Z-score was calculated by dividing the difference between the observed and shuffled means by the standard deviation of the shuffled data. The p-value is derived from the Z-score using the two-tailed Gaussian cumulative distribution function. For the comparison between categorical data (including Boolean distribution, e.g., Fig. 2G), Chi-Square test was used.

### Spike coupling

The spike coupling was defined by the baseline corrected cross-correlation (BCCC) method adapted from (*70*). First, only single units with > 1Hz average spike rate were kept after spike-sorting, thus generating at least 4000 spikes during recording sessions (average duration 6072 ± 534 sec, minimum 4071 sec). Cross-correlograms (CCG, 0.4 msec binning) were generated with the spike trains from each single unit recorded in the same session. To be qualified as a significant spike coupling, the peak in the CCG needed to exceed that from the estimated baseline. To estimate the baseline, the observed CCG was convolved with a ‘‘partially hollow’’ Gaussian kernel (*70, 71*), with a standard deviation of 10 msec, and a hollow fraction of 60%.

The probability (p) of observing equal, or more spikes from the following neuron in each bin of the observed CCG (0.8 to 10 msec) was calculated using the baseline, and the continuity-corrected Poisson distribution method (*70–72*). Only neuron pairs with p < 0.001 were considered. Furthermore, the peak in the causal direction (within the positive latency window, 0.8 to 10 msec) needed to be larger than the largest peak in the anti-causal direction (within the negative latency window, −9.2 to 0 msec).

A pair of coupled neurons was deemed congruent if the two neurons preferred the same olfactory sample and could be activated by the same task condition, as illustrated in fig. S4. A pair of coupled neurons was deemed incongruent if they preferred different olfactory samples (fig. S4). If neither of two functionally linked neurons is memory-selective, the pair is deemed non-memory (fig. S4). Out of a total of 70046 SC identified, 7674 were congruent couplings, 6299 were incongruent couplings, and 50579 were non-memory couplings.

### Relation between spike-coupling rate and distance

During the aforementioned probe localization procedure, the 3D coordinates of all neurons were acquired. The averaged coordinates of all neurons in a brain region are regarded as the region’s coordinates. The spatial distance between each pair of brain regions was then calculated as the norm of the vector difference between the two regions’ coordinates. The SC rate was calculated by dividing the number of observed SCs by the total number of possible SCs for each pair of brain regions. Using the MATLAB function *fitnlm*, the SC rate and spatial distance were then fitted with a power-law nonlinear regression model R = β1*(D^β2^) + β3, where R represented SC rate and D represented the distance between brain regions.

### CSP SVM Classifier analysis

The support vector machine classifier analysis for the CSPs (Fig. 2E and fig. S4C) was similar to the classification analysis for neuronal firing rates, with the exception that the frequency of CSPs was used in place of the neuronal firing rates, and 100 iterations of 2-fold cross-validation were performed to estimate the variance. The classification accuracy was evaluated for congruent, incongruent, and non-memory SCs in incremental numbers of 5, 10, 50, and 100 to 500 pairs in steps of 100.

### Activity chains

In the previously described SC dataset, some neurons acted as the following neuron in one SC and the leading neuron in another SC. Such a neuron thus creates a chain by connecting two SCs and three neurons. Using the same concept, one may extend these chains farther.

Conforming to the definition in SCs, a congruent chain is defined as a chain of neurons that are congruent to all other neurons in the chain. The cross-region activity chains were further classified as consistent, or inconsistent, by the wave timing of associated brain regions. In a consistent chain, the wave-timing (TCOM) of the brain region of the following neuron is either the same, or later (larger) than that of the brain region of the leading neuron for all SC pairs in the chain, whereas in an inconsistent chain, the following neuron’s wave-timing is the same or earlier (smaller). Chains formed by SCs of mixed direction (both consistent and inconsistent) were observed but excluded for further analysis.

Chained spike sequences were then searched in the coupled chains using the spike time of each associated neuron. Each spike in the chained spike sequence must adhere to the leading-following SC relationship and occur within the 10-msec time window. A chained spike sequence always begins with a spike from the first neuron, continues with one spike from each neuron in the chain, and concludes with a spike from the last neuron in the chain.

In total, we found 3,044 consistent chains, 2,021 inconsistent chains, and 2887 within-region chains. Only consistent and within-region chains were included in further analysis, unless inconsistent chains were explicitly mentioned.

### Activity loops

The coupled loops were detected using similar methods for coupled chains, with the additional requirement that a linked neuron is coupled back to the initiating neuron.

Activity loops were searched in the spikes from neurons forming coupled loops. The first and last spike in an activity loop could be from any of the neurons in the loop as long as each neuron in the loop contributes spikes in sequence (one spike in each cycle) and the first neuron is recurrently activated at least once, with no restrictions on the total number of cycles. A total of 37723 single loops were identified (923 of 3 neurons, 5119 of 4 neurons, and 31681 of 5 neurons), 782 of which were congruent.

### Nested-loops

The nested-loops are aggregated assemblies of congruent chains and loops. Each of the chains or loops in a nested-loop contains at least one neuron that is also associated with other chains or loops in the assembly. Continuous activation of a nested-loop was defined as the summation of temporally overlapped chains or loops sequences of the neurons in the assembly. Multiple chains or loops may be activated by independently initiating spikes in a continuous activation of a nested-loop. For the cross-session statistics, due to the congruent requirement, each session is further grouped into 4 conditions based on sample and delay duration. Among the 182 session-conditions in which at least one congruent motif was observed, nested-loops were found in 140 session-conditions.

For the study of the effects of varied delay in connecting chains and loops in nested-loops (fig. S9A), instead of the overlapping-rule, the activity chains and loops were allowed to connect by brief gaps of 5 ms, 10 ms, or 20 ms.

### Generalized linear model for predicting behavior responses

In order to determine the effect of previous trials on behavior, responses to each trial across all sessions were collected with the sample and test odor of the current trial, as well as sample odors from three previous trials. All lick responses were assigned a value of 1, while non-lick responses were assigned a value of 0. Similarly, all trials with samples matched to the current test were assigned a value of 1; otherwise, a value of 0 was assigned. A generalized linear model was constructed where 4 weighted trials and a constant would predict the response (i.e., R = β_1_M_-3_ + β_2_M_-2_ + β_3_M_-1_ + β_4_M_0_ + β_5_, where R represent response and Ms represented the sample-test matching relationship among the trials) using the MATLAB *fitglm* function, with the distribution set to “binomial” and link function set to “identity”. The β values were illustrated in figure S7B.

### Analysis to remove specific components from nested loops

For the analysis of changes following removal of specific components from the nested loops, the experimentally observed nested chains comprised of at least two component motifs (chains or loops) were used as reference. First, all the SCs, activity chains, single loops, and HIP region neurons in each of these nested loops were labeled. Second, for each template and each type of the components, a subgroup of a given label was chosen and excluded, and the nested loops constructed from the remaining components were compared against the reference. Such remove-rebuild-compare process was systematically repeated up to 100 times in each step, if the total number of potential combinations exceeded 100. Otherwise, all the possible combinations for removal were examined once. When specific SCs, chains, or loops were being considered for removal, if a neuron or SC was present in both the pending-removal motif and other preserved motifs, the neuron or SC was not removed. The removal of HIP neurons occurs after the formation of nested-loops, so a neuron selected for removal is eliminated regardless of the number of motifs in which it participated.

### Data and code availability

The computer programs used in this study have been deposited publicly at https://gitee.com/XiaoxingZhang/pixels and https://github.com/StringXD/SmartTrack-Histology.

The data have been deposited publicly at https://datadryad.org/stash/share/Z7wUPpxCmJWM621LgU4TjNxakIlV_CFl-op1nc4n9lo.

## Supplementary Text

### Behavior performance

The well-trained criterion is performance correct rate ≥ 75% in 40 consecutive trials, which is consistent with previous studies using a similar behavioral task design(*14, 15*). For the analysis of correct behavior, only the hit and correct-rejection trials within well-trained windows were included. For the analysis of incorrect behavior, error trials in the whole session (both within and outside the well-trained window) were included.

### AUROC

If the neurons of a given activity pattern are important for WM task performance, one would expect failure of WM encoding in these neurons in the error trials. Thus, we separately examined the coding ability of transient and sustained neurons in the correct and error trials, with the focus on the delay period. The distribution of activity modulation in transient neurons showed clear separation for preferred and nonpreferred odor in the correct trials, but not in error trials (fig. S2C), thus was tightly related with behavioral performance. This is also revealed by the value of area under receiver operating characteristic curve (AUROC; 0.99 for correct trials, and 0.65 for error trials, fig. S2D). The activity of the sustained odor neurons could significantly encode the preferred odor in both correct and error trials (1.00 for correct trials, and 0.86 for error trials, fig. S2D). Thus, the activity of transient neurons was more correlated with behavioral performance.

**Movie S1.** Performance of a DPA task in mice.

---

**Fig. S1.**
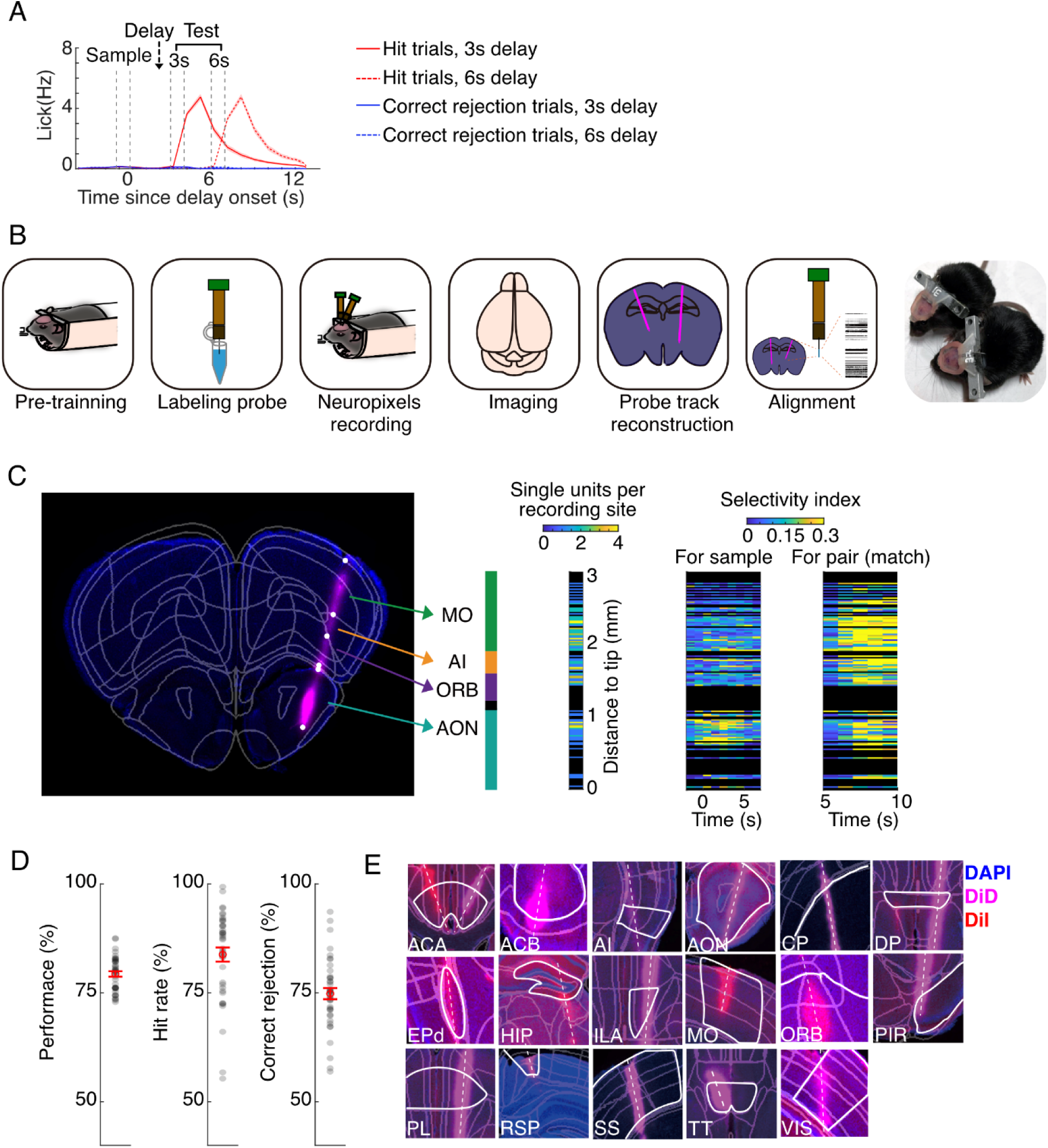
Paradigm of Neuropixels-probe recordings in behaving mice. (**A**) Averaged lick rates for the corresponding session in Figure 1H. (**B**) Schematic diagram and head-bar implementation for Neuropixels probe recording in awake, behaving mice. (**C**) Alignment of recording sites and registration of single units to brain regions. Left, fluorescence labeling of an example recording track. Middle, regions registered according to Allen Brain Atlas. Right, reference electrophysiology features for the corresponding sites, including single-unit density, averaged odor selectivity-index, and averaged pair/non-pair selectivity-index. For clarity, only data from 6 sec-delay trials were shown. (**D**) Behavioral performance (77.1% ± 0.9%), hit (80.2% ± 2.0%), and correct rejection (74.1% ± 1.5%) rates of the mice in the ODPA task. Black dots represent data from 40 individual mice, while red markers represent mean ± S.E.M. (**E**) Example histological images for recording tracks. Contours marked in white indicate brain region outline, with the region name listed in the left-bottom corner of each plot.

**Fig. S2.**
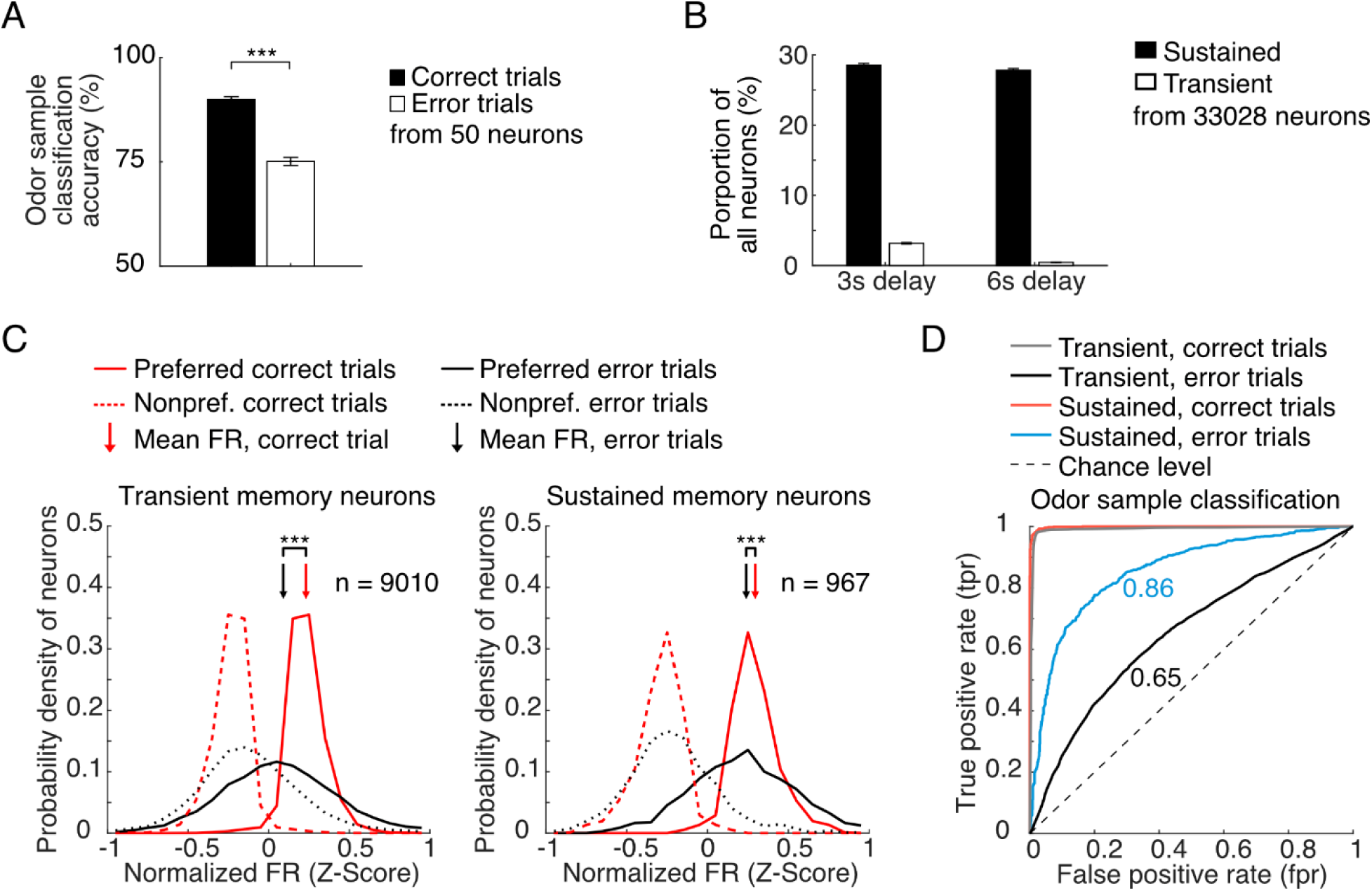
Transient patterns of memory-selective activity associated with WM. (**A**) Classification accuracy using the delay-period firing rate of 50 neurons, in cross-validated SVM classifier analysis that decodes for the odor sample, in the correct and error trials. ***, p < 0.001, Chi-Square test. (**B**) Proportion of sustained- and transient-selective neurons across all sessions in the 3-sec or 6-sec delay period following sample offset. Error bars represent S.E.M. (**C**) Probability distribution of the normalized firing rates in the preferred and non-preferred trials, for the transient (left) and sustained (right) memory neurons, in the correct and error trials. (**D**) AUROC for transient or sustained memory neurons in correct and error trials, derived from (C).

**Fig. S3.**
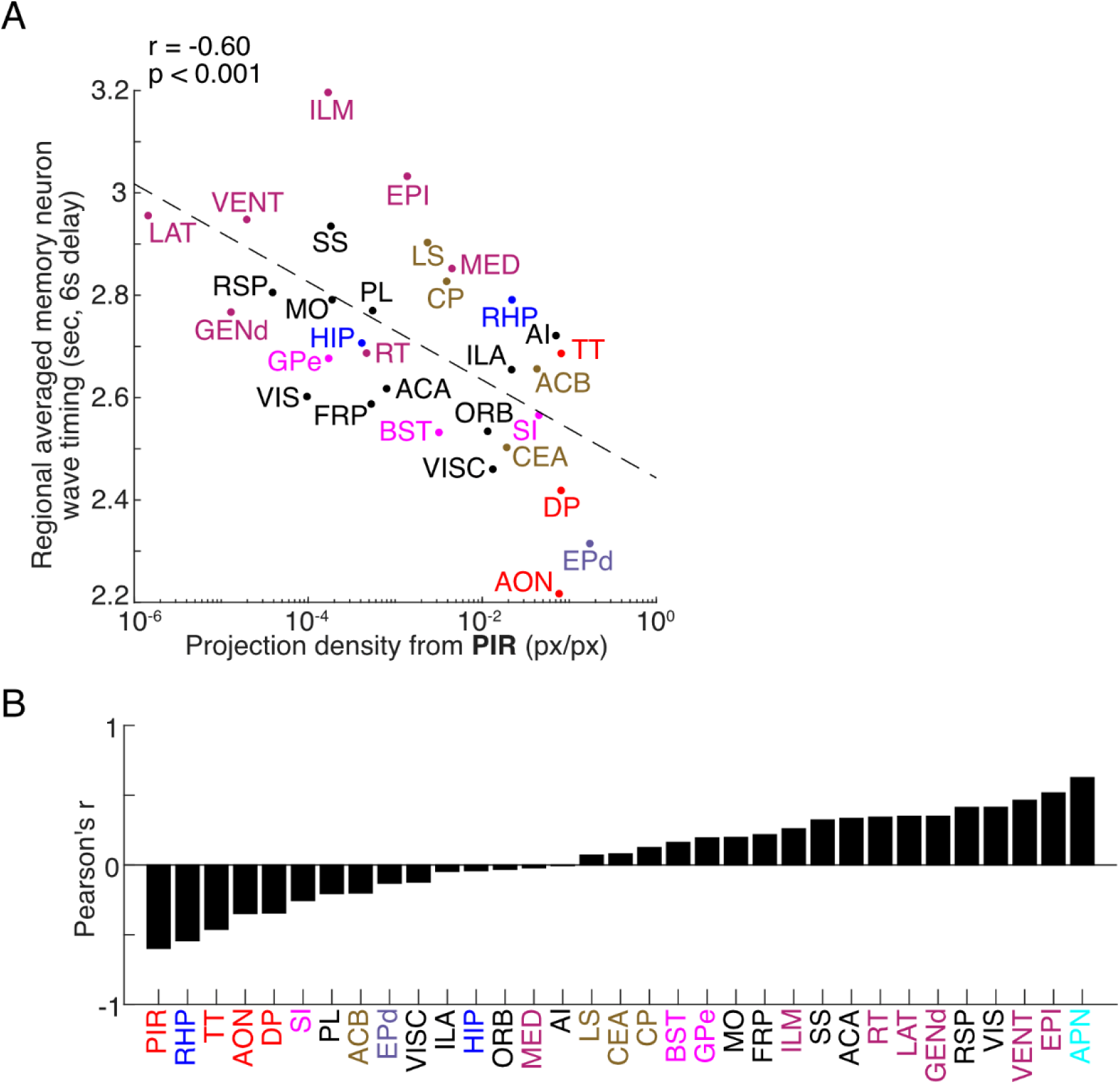
Relationship between regional wave timing and anatomy. (**A**) Correlation between regional averaged wave timing of memory neurons in 6 sec-delay trials, and projection density from PIR to each of these regions using data from the Allen Brain Atlas. (**B**) Distribution of correlation coefficients between regional averaged wave timing of memory neurons and projection density for the recorded regions as in (A).

**Fig. S4.**
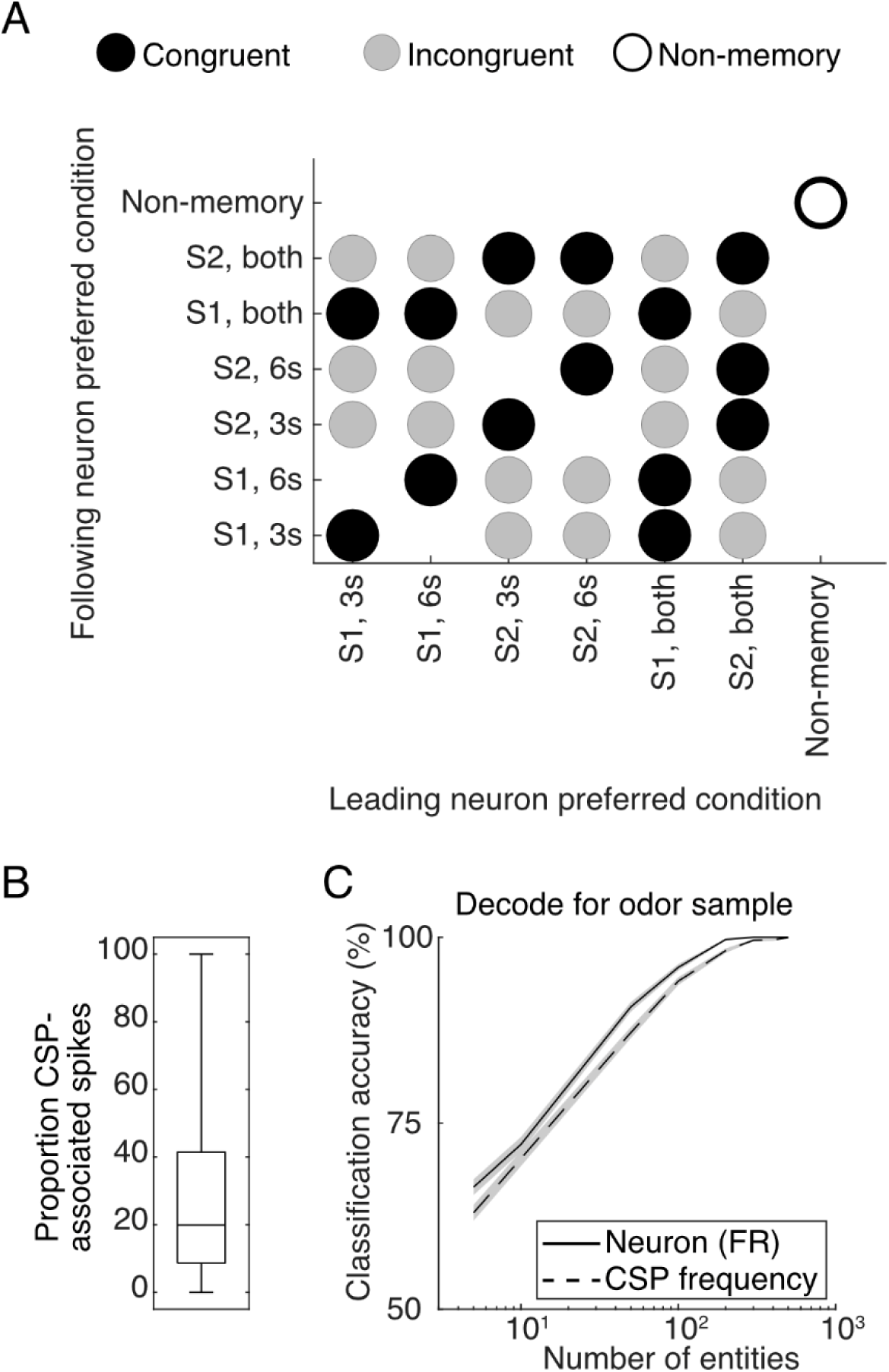
Congruent, incongruent, and non-memory relationships in coupled neuron pairs. (**A**) A detailed illustration of the inter-neuron relationship in terms of the memory specificity of the leading and following neurons. Unmarked pairs (e.g., between non-memory neurons and memory neurons) were excluded from further study. **(B)** Proportion of spikes that are associated with congruent CSPs among memory neurons. (**C**) Classification accuracy for sample odor, using the CSP frequency of varied number of memory-neuron pairs (each pair was counted as one entity), or firing rates of a varied number of memory neurons.

**Fig. S5.**
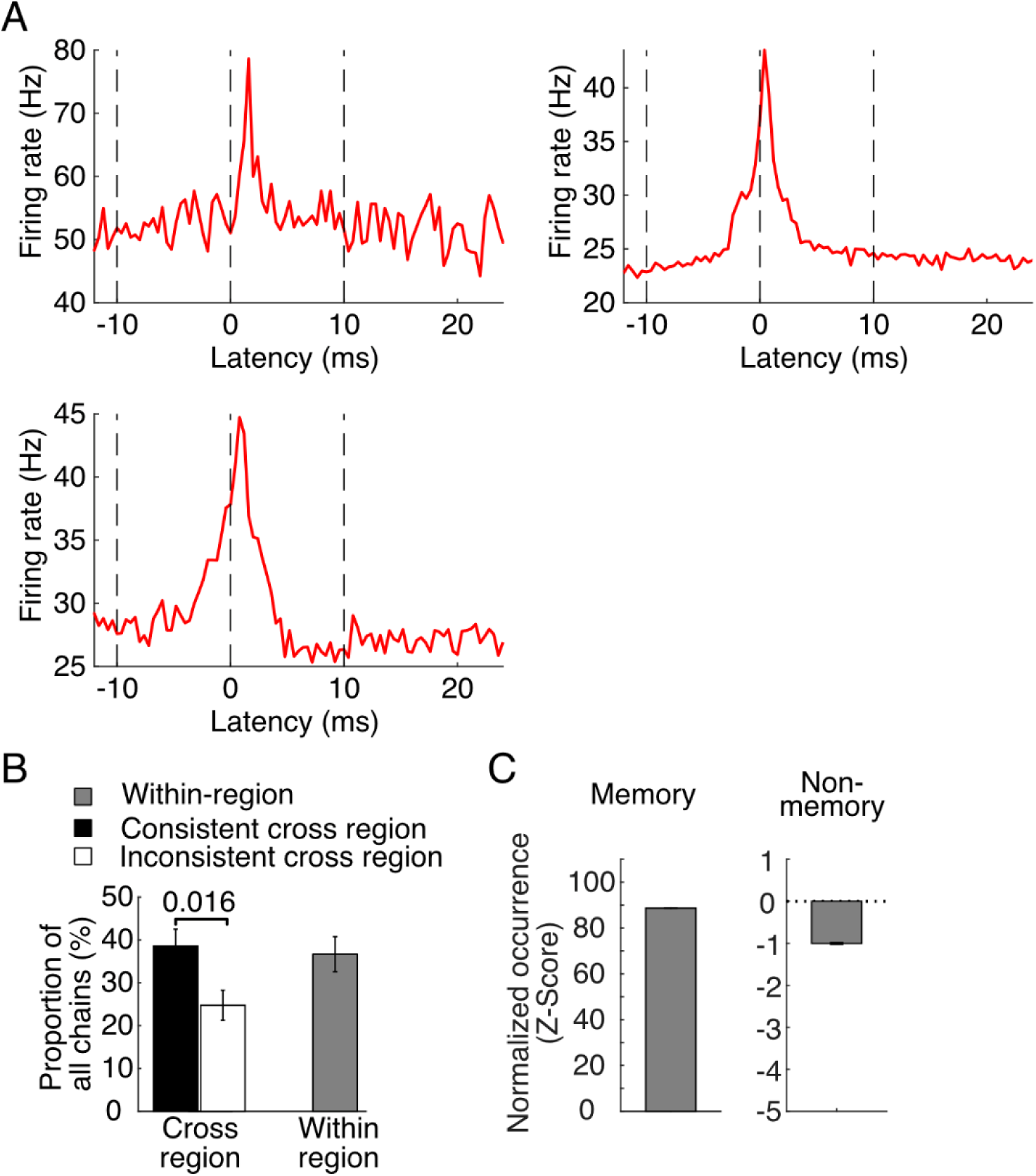
Memory-selective activity chains and loops. **(A)** Spike cross-correlograms of the series of neuronal pairs in the example activity chain in Figure 2H. (**B**) Proportion of SC-connected neuronal chains in the consistent or inconsistent direction, or within-region, for the memory neurons. P-value from rank-sum test across sessions. (**C**) Shuffle-normalized ratio (Z-score) of within-region chains.

**Fig. S6.**
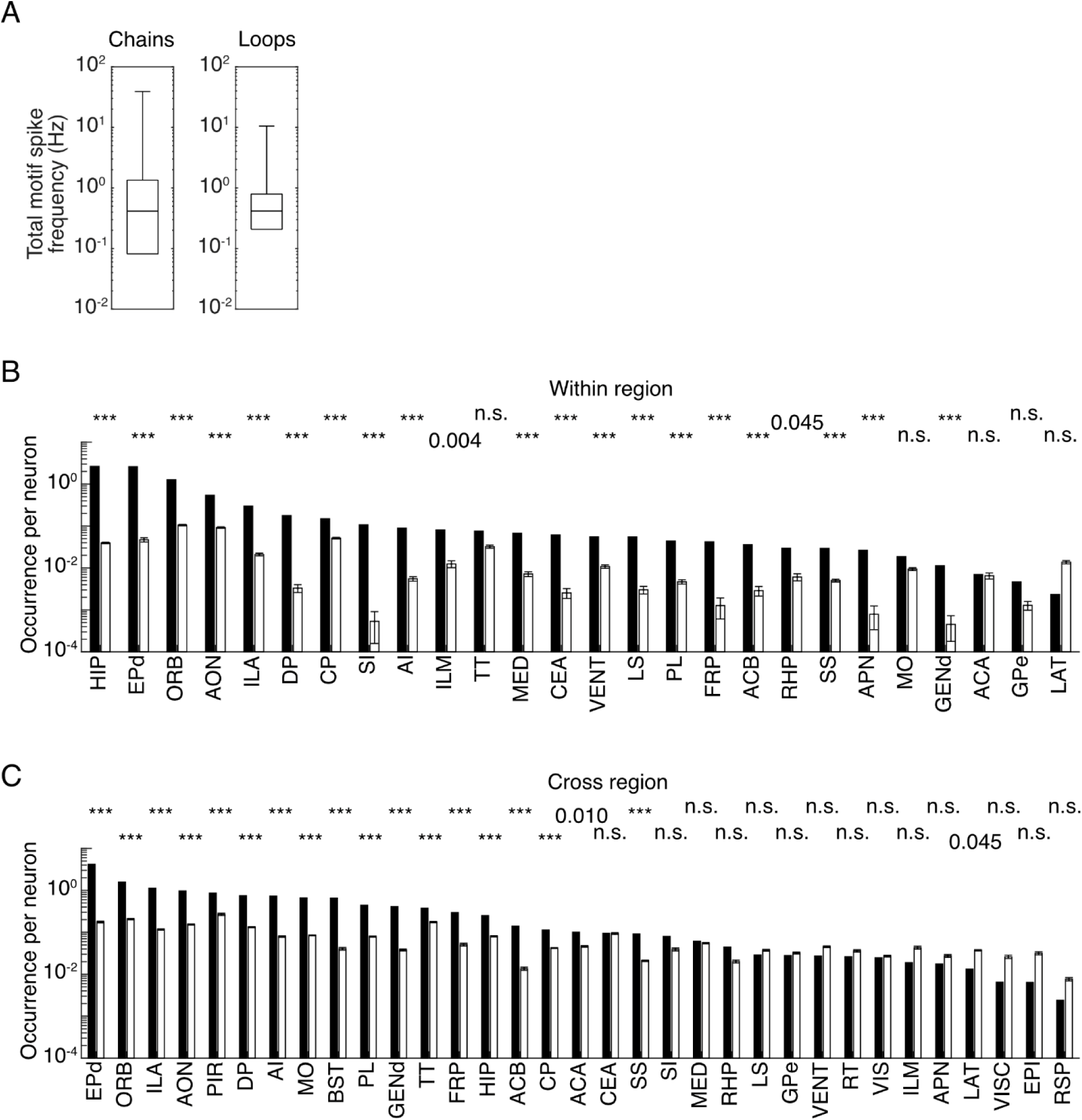
Memory-selective activity chains, and single loops. **(A)** Spike frequency of observed activity chains (left) and single loops (right) throughout the delay duration. Statistics from individual sessions, whiskers represented the full range. (**B**) Normalized occurrence of neurons in the within-region activity chains (divided by the total number of record neurons in the region, across all regions with at least one neuron in chains). Black bar represented observed data, white bar represented shuffled data (from control datasets in which the number of neurons and SCs were the same, but the couplings between neurons were random). ***, p< 0.001, n.s., not significant, probability from 100 shuffled repeats. (**C**) Similar to (B), but for cross-region activity chains.

**Fig. S7.**
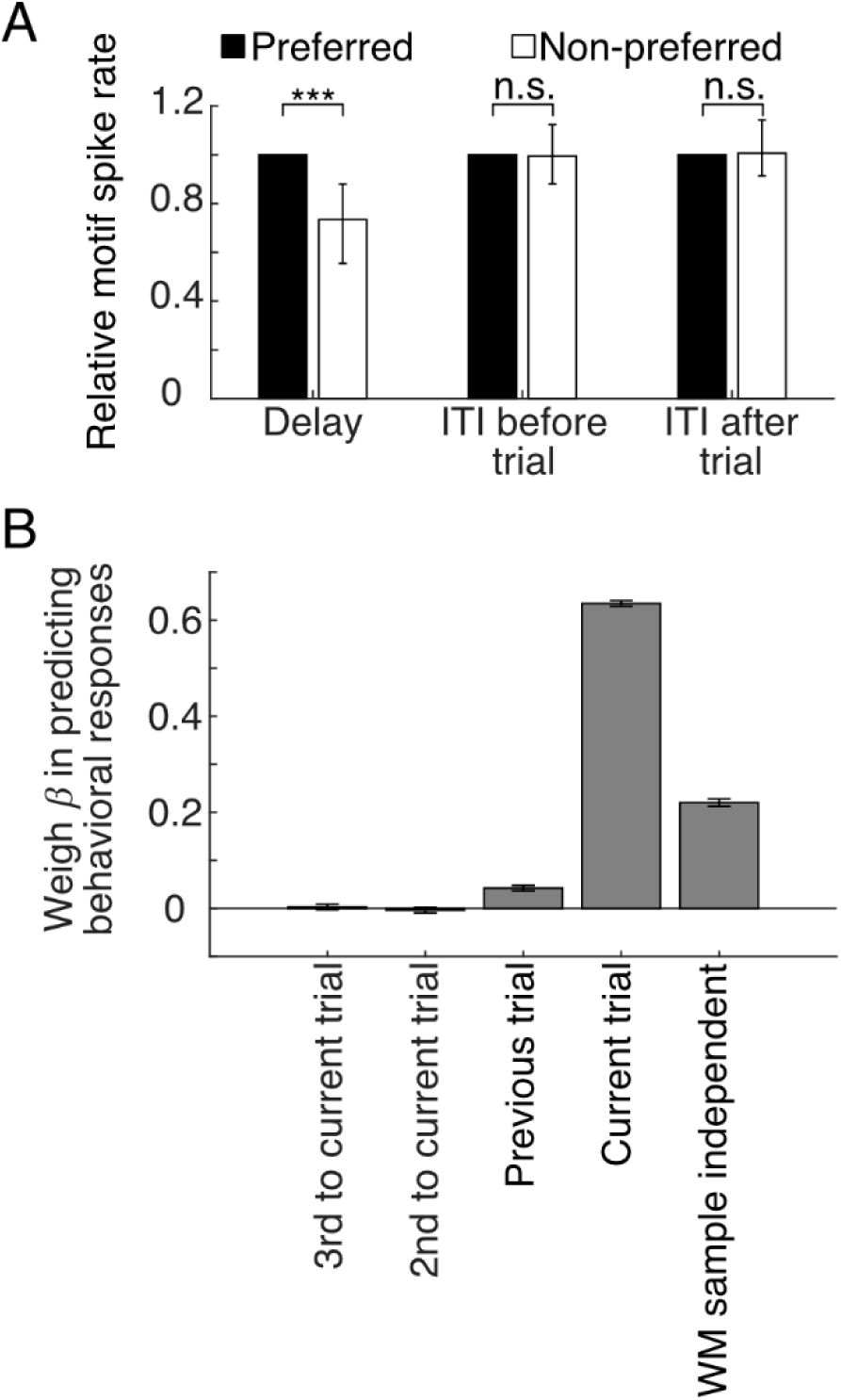
Replays of WM-associated activity chains and loops. **(A)** Reduced motif activity in the non-preferred trials during the delay period, and similar replayed motif activity in the ITI periods before and after a trial. Bars and error bars represent the median and 95% C.I., ***, p < 0.001, n.s., not significant, signed-rank test, as above. (**B**) Weigh (*β*) in a fitted generalized linear model predicting behavioral responses (lick / no lick). The response is predicted from the pair relationship between the test odor in the current trial, and the sample odor in the current and three previous trials. Higher *β* represents higher power in prediction. Error bars represent standard error.

**Fig. S8.**
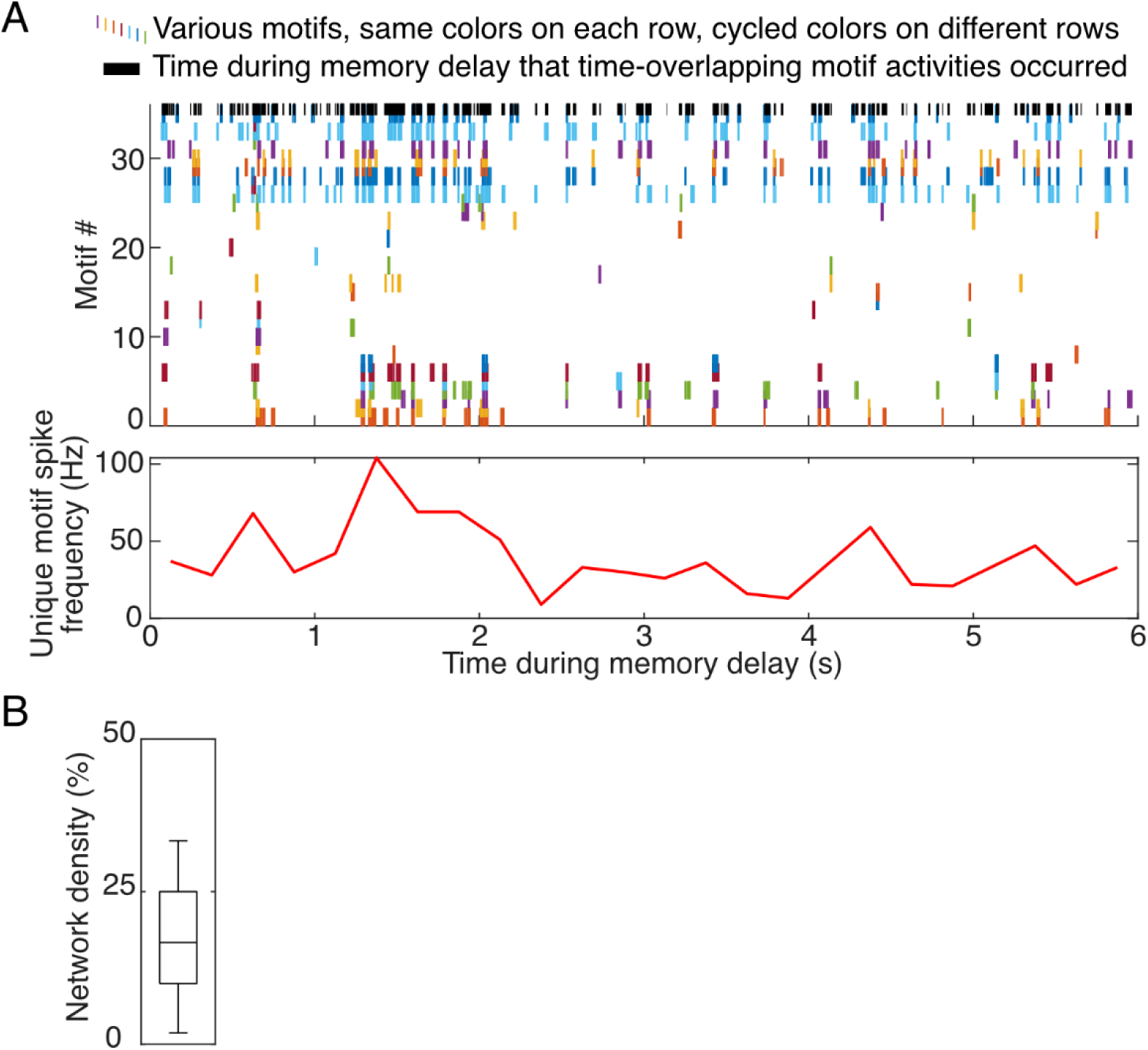
Nested loops and hierarchical organization of cross-region sequential spiking. (**A**) Raster plot of spiking of chains and single loops involved in the nested loop in Figure 4D. Above: colored bars represent sequential alternative activation of various motifs at a 10-msec resolution. Black bars represented parts of the WM delay covered by overlapping motif activities. Below: combined motif-associated spike frequency during this period. (**B**) Graph network connection density (defined by the number of actual SC divided by that of all possible SC) within the observed nested loops. Whiskers represented full range.

**Fig. S9.**
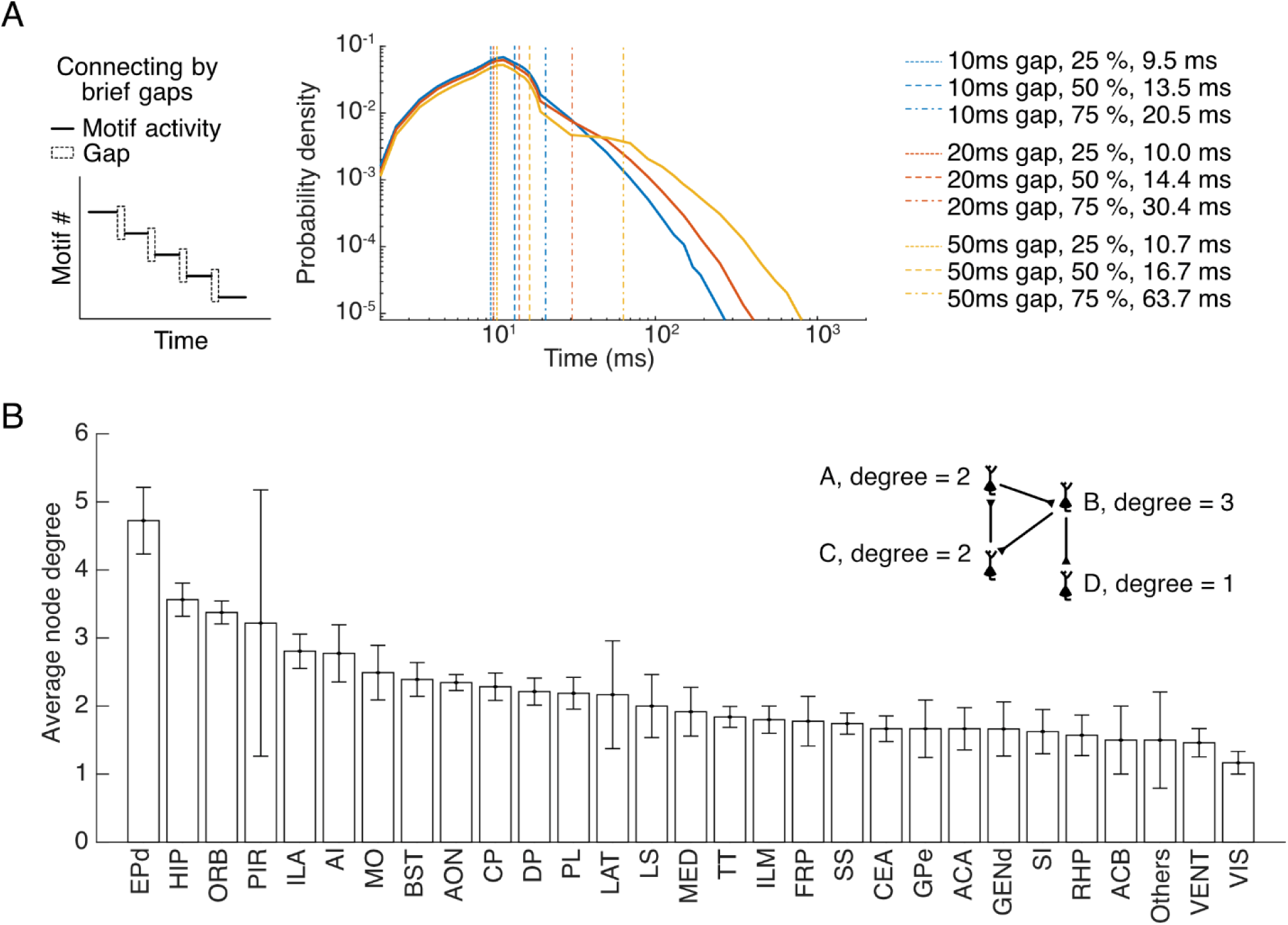
Replays of WM-associated activity chains and loops. (**A**) Duration distribution for activity-sequences, defined as consecutive activities of constituent chains or loops in the nested-loops, connected by brief gaps of varied maximum allowed width. (**B**) Average node degree in various regions of the brain. Regions that contributed more than five neurons in the nested loops were estimated individually, while the remaining regions were grouped together as “others”. Inset: Method for computing node degree in nested loops (shown as numbers following neuron-labels A-D), which indicated the number of input and output SCs connecting each neuron to other neurons.

**Fig. S10.**
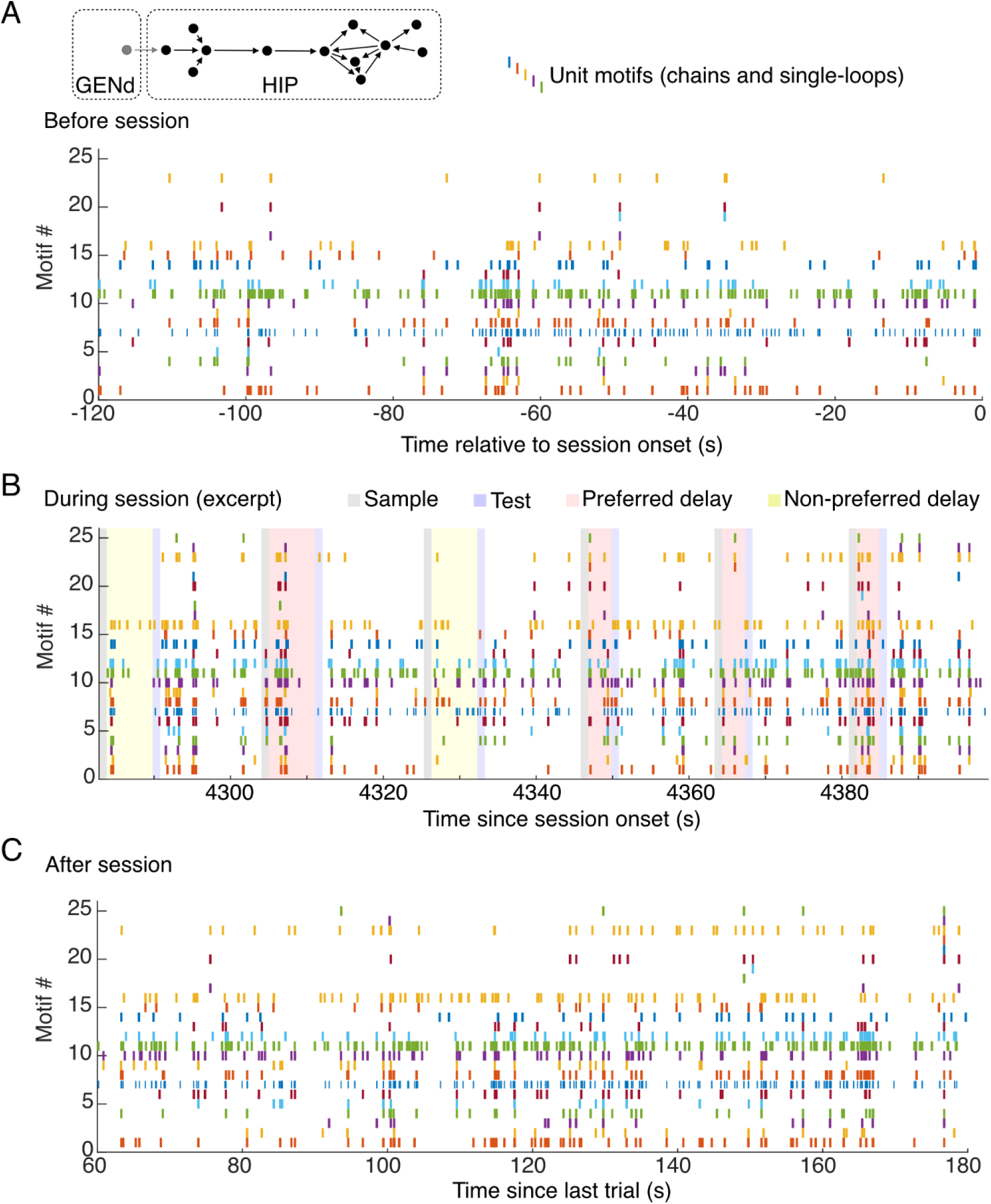
Replays of nested loops. (**A**) Unit motifs of a nested-loops structure spontaneously replayed before the start of the task execution. Inset: diagram of the nested-loops structure, which was comprised of 12 HIP neurons and one GENd neuron that formed 25 child-motifs. (**B**) Occurrence of the same unit motifs as in (A), during and between the delay periods. Color shades labeled above: corresponding time periods. (**C**) as in (A), after the end of task execution.

**Table S1.** Memory neurons across brain regions.

| Index | Abbreviation | Name | All neurons | Memory neurons |
| --- | --- | --- | --- | --- |
| 1 | CP | Caudoputamen | 3512 | 902 |
| 2 | MO | Somatomotor areas | 2899 | 533 |
| 3 | ORB | Orbital area | 2491 | 1091 |
| 4 | SS | Somatosensory areas | 2108 | 380 |
| 5 | HIP | Hippocampal region | 1869 | 670 |
| 6 | LAT | Lateral group of the dorsal thalamus | 1283 | 273 |
| 7 | AON | Anterior olfactory nucleus | 988 | 583 |
| 8 | PL | Prelimbic area | 866 | 294 |
| 9 | AI | Agranular insular area | 777 | 235 |
| 10 | VENT | Ventral group of the dorsal thalamus | 703 | 155 |
| 11 | GPe | Globus pallidus external segment | 644 | 145 |
| 12 | ILA | Infralimbic area | 594 | 283 |
| 13 | MED | Medial group of the dorsal thalamus | 582 | 211 |
| 14 | VIS | Visual areas | 488 | 86 |
| 15 | TT | Taenia tecta | 435 | 290 |
| 16 | ACA | Anterior cingulate area | 428 | 139 |
| 17 | RSP | Retrosplenial area | 417 | 68 |
| 18 | LS | Lateral septal nucleus | 416 | 132 |
| 19 | DP | Dorsal peduncular area | 344 | 176 |
| 20 | RT | Reticular nucleus of the thalamus | 304 | 78 |
| 21 | CEA | Central amygdalar nucleus | 294 | 114 |
| 22 | GENd | Geniculate group dorsal thalamus | 265 | 52 |
| 23 | ACB | Nucleus accumbens | 250 | 84 |
| 24 | ILM | Intralaminar nuclei of the dorsal thalamus | 211 | 60 |
| 25 | RHP | Retrohippocampal region | 202 | 77 |
| 26 | GPi | Globus pallidus internal segment | 180 | 52 |
| 27 | EPd | Endopiriform nucleus dorsal part | 166 | 107 |
| 28 | BST | Bed nuclei of the stria terminalis | 159 | 46 |
| 29 | EPI | Epithalamus | 156 | 41 |
| 30 | VISC | Visceral area | 154 | 45 |
| 31 | FRP | Frontal pole cerebral cortex | 142 | 40 |
| 32 | APN | Anterior pretectal nucleus | 114 | 29 |
| 33 | PIR | Piriform area | 114 | 52 |
| 34 | SI | Substantia innominata | 112 | 45 |
| 35 | ATN | Anterior group of the dorsal thalamus | 86 | 18 |
| 36 | AUD | Auditory areas | 81 | 16 |
| 37 | LPO | Lateral preoptic area | 57 | 20 |
| 38 | FS | Fundus of striatum | 47 | 16 |
| 39 | RPF | Retroparafascicular nucleus | 42 | 8 |
| 40 | GU | Gustatory areas | 39 | 3 |
| 41 | NPC | Nucleus of the posterior commissure | 32 | 6 |
| 42 | AAA | Anterior amygdalar area | 29 | 11 |
| 43 | SH | Septohippocampal nucleus | 29 | 20 |
| 44 | EPv | Endopiriform nucleus ventral part | 23 | 16 |

**Table S1. Memory neurons across brain regions.**
| Index | Abbreviation | Name | All neurons | Memory neurons |
| --- | --- | --- | --- | --- |
| 45 | BMAa | Basomedial amygdalar nucleus anterior part | 22 | 5 |
| 46 | MEA | Medial amygdalar nucleus | 21 | 5 |
| 47 | IA | Intercalated amygdalar nucleus | 20 | 11 |
| 48 | PTLp | Posterior parietal association areas | 20 | 9 |
| 49 | SCig | Superior colliculus motor related intermediate gray layer | 17 | 4 |
| 50 | ECT | Ectorhinal area | 15 | 4 |
| 51 | GENv | Geniculate group ventral thalamus | 13 | 1 |
| 52 | MOB | Main olfactory bulb | 13 | 9 |
| 53 | PRC | Precommissural nucleus | 13 | 2 |
| 54 | TEa | Temporal association areas | 13 | 3 |
| 55 | PS | Parastrial nucleus | 12 | 1 |
| 56 | PP | Peripeduncular nucleus | 10 | 1 |
| 57 | OP | Olivary pretectal nucleus | 9 | 0 |
| 58 | BMAp | Basomedial amygdalar nucleus posterior part | 7 | 4 |
| 59 | MA | Magnocellular nucleus | 4 | 2 |
| 60 | NLOT | Nucleus of the lateral olfactory tract | 4 | 1 |
| 61 | BAC | Bed nucleus of the anterior commissure | 1 | 1 |
| 62 | MPO | Medial preoptic area | 1 | 0 |

